# Integrative scRNA-seq and spatial transcriptomics uncovers distinct macrophage-fibroblast cross-talk in human hip synovium between patients with femoroacetabular impingement and osteoarthritis

**DOI:** 10.1101/2024.12.09.627061

**Authors:** Gulzada Kulzhanova, Alexis Klee, Mina Botros, Victoria L. Hansen, John Reuter, Eliya Tazreena Tashbib, Eloise Fadial, Benjamin F. Ricciardi, Brian Giordano, Chia-Lung Wu

## Abstract

Femoroacetabular impingement (FAI) and synovitis have been recognized as essential factors for developing osteoarthritis (OA) in the hip joints. However, little is known about altered synovial cellular compositions, their associated transcriptomic profiles, and cell-cell interactions between patients with FAI and hip OA. In the current study, by using integrative single-cell RNA sequencing (scRNA-seq) and spatial transcriptomics (Spatial-seq), we identified the molecular mechanisms by which synovial cells promote hip OA pathogenesis from FAI. Compared to FAI synovium, epiregulin (EREG)-enriched lining fibroblast-like synoviocytes (FLS) were significantly increased in the hip OA synovium. These *EREG^+^* FLS are pro-inflammatory due to their high expression of *CXCL1*, *IL8 (CXCL8)*, and *MMPs*. Furthermore, pseudotime analysis predicts that *EREG^+^* FLS are potentially derived from *DPP4^+^PI16^+^* sublining FLS. Importantly, analysis of cell-cell interactions indicates that fibroblast growth factor 2 (FGF2) secreted from *COL1A1^+^IGFBP5^+^* fibrotic macrophages may signal through syndecan 4 (SDC4) expressed by *EREG^+^* lining FLS, inducing the expression of IL6, IL8, MMP1, and PTGS2. The GO term analysis of activated genes downstream of FGF2-SDC4 signaling revealed that biological processes associated with inflammation and angiogenesis were upregulated in hip OA, while mechanical stimulus and skeletal muscle differentiation were dominant in FAI. Moreover, we also found that *EREG^+^CCL20^+^MMP3^hi^* lining FLS as well as most MΦ and monocyte populations are unique to hip OA patients when compared to knee OA and RA patients. The findings of this study offer a groundwork in tailoring novel targets and therapies for FAI and hip OA patients.

## Introduction

Cam-type femoroacetabular impingement (FAI) is a pathomorphological condition whereby the abnormal osseous protrusion at the femoral head-neck junction impinges on the acetabular rim leading to acetabular labral injury [1], and aberrant loading on the femoral articular cartilage and subchondral bone [1, 2]. More importantly, it has been shown that FAI can lead to aberrant loading on the femoral articular cartilage and subchondral bone, creating an unstable biomechanical environment [1, 2]. FAI has been considered as a primary cause of hip and groin pain in young individuals aged 15-50 years, especially affecting individuals involved in high-intensity activities [1, 3–6]. Even though the causal relation between FAI and hip osteoarthritis (OA) has not been fully established, Cam-type FAI is highly associated with hip OA development [1, 3, 6–8]. For example, patients with Cam-type deformity may have a higher risk of developing hip OA within 5 years [4]. Additionally, the levels of inflammatory cytokines such as IL1-𝛽 in late-stage FAI synovial tissues have been reported to be elevated compared to early-stage FAI [3]. The tandem of abnormal contact forces and synovial inflammation in Cam-type FAI joints may contribute to cartilage delamination and labral pathology, gradually leading to development of hip OA [1, 2, 4]. Thus, FAI has been considered a unique early-phase hip OA model for studying regulators implicated in hip OA onset and progression.

OA is characterized by the progressive degeneration of articular cartilage, subchondral bone sclerosis, and synovial inflammation [9, 10]. Due to the increasing elderly population, the incidence of OA is projected to double by 2045 [11]. Currently, hip OA affects approximately a quarter of humans worldwide by age 85 [12]. While a plethora of factors including aging, obesity, genetic background, and abnormal biomechanical loading may contribute to hip OA development, a growing body of evidence also suggests that a low-grade inflammation of synovium (i.e., aberrant biochemical environment) could be a major driving force [9, 13, 14].

Healthy synoviocytes maintain joint metabolic homeostasis by nourishing chondrocytes and removing the degraded cartilage extracellular matrix [10]. However, in pathological scenarios, abnormal behavior of synoviocytes may lead to enhanced proliferation of fibroblast-like synoviocytes (FLS), elevated angiogenesis, and increased recruitment of immune cells, consequently leading to synovial hyperplasia and inflammation [9, 10]. It is well-documented that inflammatory cytokines derived from the synovial cells can promote OA in knee joints [15]. Recently, the cellular composition of synovium in rheumatoid arthritis (RA) and OA within the knee joints has been extensively characterized by means of next-generation sequencing and bioinformatics [15–17]. However, the unique cytokine profiles of the synovial fluid between knee and hip OA joints suggest that hip OA is immunologically distinct from knee OA [15]. Thus, there is a significant knowledge gap regarding the cellular composition and their associated transcriptomics as well as cell-cell interactions in the synovium during hip OA progression. Moreover, the molecular mechanisms by which synovial cells contribute to FAI-to-hip-OA progression remain elusive. In this study, we hypothesize that synovial cells from FAI and hip OA patients exhibit distinct transcriptomic profiles and cell-cell interactions. We aim to identify the molecular mechanisms underlying hip OA pathogenesis from FAI by integrating innovative single-cell RNA sequencing (scRNA-seq) and spatial transcriptomics (Spatial-seq) approaches. Furthermore, elucidating signaling pathways and their downstream activated genes in the synovial cells will allow us to identify target genes for therapeutic intervention for FAI and hip OA patients.

## Methods and Materials

### Synovium sample harvest

Ethical approval was granted by the Research Subjects Review Board at the University of Rochester Medical Center (URMC), and informed written consents were obtained from all patients participating in this study prior to surgery. Synovium tissues were harvested from the regions with the most severely damaged cartilage from the femoral head-neck junction of patients with Cam-type FAI and advanced hip OA (secondary to FAI) undergoing hip arthroscopy and total hip arthroplasty surgeries, respectively. Details on sex, age, ethnicity, and body mass index (BMI) were listed in **Supplemental Table 1**. FAI and hip OA synovium samples were collected in a tissue collection medium (DMEM/F-12, GlutaMAX™ supplement, #10-565-018, Gibco™, with 1% Penicillin-Streptomycin, #15140122, Gibco™, and 10% Fetal Bovine Serum, #F0926-100, Sigma-Aldrich) for single-cell RNA-sequencing (scRNA-seq), spatial transcriptomics, and/or immunofluorescent staining (**Fig. 1**).

**Fig. 1.**
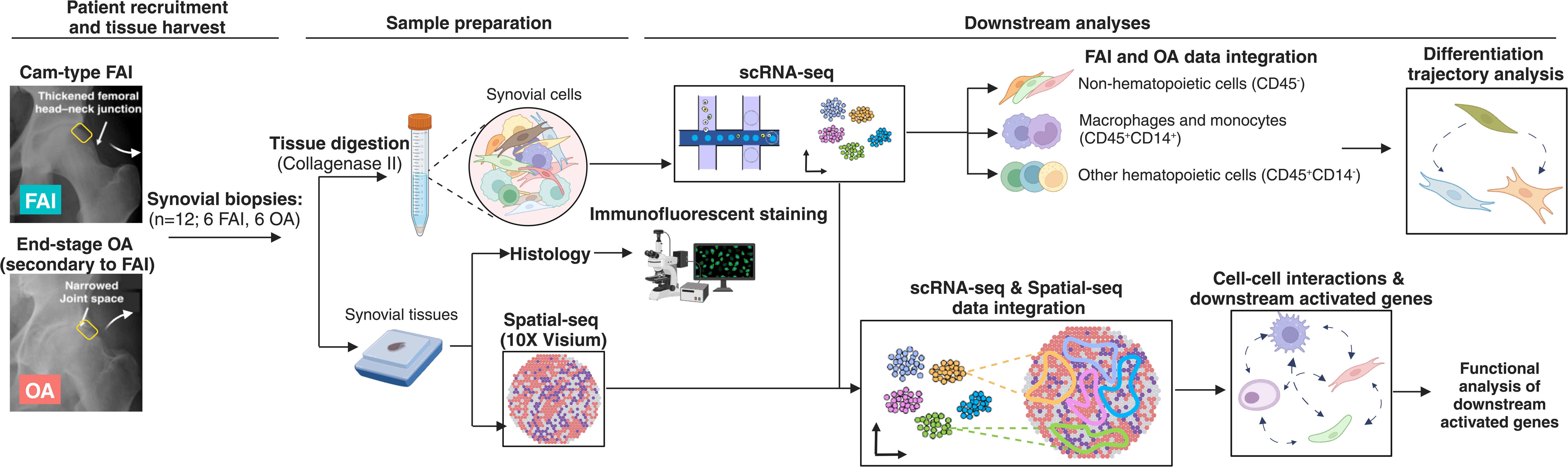
An overview of experimental workflow. Twelve synovial samples (n = 3/disease condition/sex) were harvested from FAI and OA hip joints, digested by collagenase type II, and profiled using scRNA-seq. Cells were grouped based on general markers CD45 and CD14: (1) *CD45^-^*non-hematopoietic cells, (2) *CD45^+^CD14^+^*myeloid cells, and (3) *CD45^+^CD14^-^* other hematopoietic cells. Non-hematopoietic and myeloid cells from all samples were integrated using Seurat R package. The identified populations were annotated using conventional synovial markers and based on prior literature. The top 50 genes were utilized for GO term functional analysis of each cell cluster using DAVID website. Furthermore, pseudotemporal ordering and TFs governing differentiation trajectory analysis were performed using Monocle 3 and RcisTarget R packages, respectively. Next, we integrated scRNA-seq datasets with Spatial-seq datasets to reveal spatial localization of the synovial cells. The populations closely adjacent to each other were chosen to investigate the cell-cell crosstalk as well as downstream activated genes using MultiNicheNet analysis.

### Primary synovial tissue digestion and preparation

Twelve human FAI and hip OA synovium samples (3 samples per disease condition per sex) were harvested for scRNA-seq analysis. The FAI synovium samples were minced by hand and digested in 0.3% collagenase medium (DMEM/F-12, GlutaMAX™ supplement, #10-565-018, Gibco™ collagenase II, #LS004177, Worthington Biochemical Corporation, and 10% Fetal Bovine Serum, #F0926-100, Sigma-Aldrich) for three hours at 37°C while shaking continuously. The samples were thoroughly vortexed every hour. The OA synovium samples were digested in 0.4% collagenase medium (DMEM/F-12, GlutaMAX™ supplement, #10-565-018, Gibco™ collagenase II, #LS004177, Worthington Biochemical Corporation, and 10% Fetal Bovine Serum, #F0926-100, Sigma-Aldrich) at 37°C overnight. Cells were then resuspended in ACK lysing buffer (#A1049201, Gibco) for 1-2 minutes to lyse red blood cells. Afterwards, cells were filtered through 70μm (#130-110-916, Miltenyi Biotec) and 30 μm MACS® SmartStrainers (#130-110-915, Miltenyi Biotec). The cell suspension was later frozen down in freezing medium (DMEM/F-12, GlutaMAX™ supplement, #10-565-018, Gibco™ with 10% Pierce™ Dimethylsulfoxide (DMSO), LC-MS Grade, #85190, ThermoFischer, and 80% Fetal Bovine Serum, #F0926-100, Sigma-Aldrich) until submitted for scRNA-seq.

### Single cell RNA sequencing

Samples with greater than 70% viability were submitted to UR Genomics Research Center for scRNA-Seq library preparation using Chromium Next GEM Single Cell 3′ GEM, Library and Gel Bead Kit v3.1 (10x Genomics), per the manufacturer’s recommendations, as summarized below. Samples were loaded on a Chromium Single-Cell Instrument (10x Genomics, Pleasanton, CA, USA) to generate single-cell GEMs (Gel Bead-in-Emulsions). GEM reverse transcription (GEM-RT) was performed to produce a barcoded, full-length cDNA from poly-adenylated mRNA. After incubation, GEMs were broken, the pooled GEM-RT reaction mixtures were recovered, and cDNA was purified with silane magnetic beads (DynaBeads MyOne Silane Beads, ThermoFisher Scientific). The purified cDNA was further amplified by PCR to generate sufficient material for library construction. Enzymatic fragmentation and size selection were used to optimize the cDNA amplicon size and indexed sequencing libraries were constructed by end repair, A-tailing, adaptor ligation, and PCR. The final libraries contain the P5 and P7 priming sites used in Illumina bridge amplification. Sequence data was generated using the Illumina NovaSeq 6000 platform.

### Spatial transcriptomics (Spatial-seq)

To reveal spatial locations of various synovial cells from FAI and hip OA patients, we performed 10X Visium spatial-transcriptomics (Spatial-seq). Human FAI (n = 4) and hip OA (n = 6) synovial tissues were isolated and placed in collection medium (DMEM/F-12, GlutaMAX™ supplement, #10-565-018, Gibco™, with 1% Penicillin-Streptomycin (10,000 U/mL), #15140122, Gibco™, and 10% Fetal Bovine Serum, #F0926-100, Sigma-Aldrich). Harvested synovial tissues were then transferred to 10% neutral buffered formalin solution. Upon 24-hour fixation at room temperature and paraffin embedding, FAI and hip OA synovial samples were sectioned at 8 μm in RNase-free environment (#7003 ThermoFischer) and placed on the individual spatially barcoded capture areas of a 10X Genomics Visium Gene Expression Slide (10X Genomics, Pleasanton, CA). Following tissue placement, slide drying, and overnight incubation in a desiccator, the slide was deparaffinized, H&E stained, and imaged using a VS120 Slide Scanner (Olympic) at 20X magnification. Sequence ready libraries were then constructed according to the manufacturer’s instructions (10X Genomics, CG000407). Final libraries were sequenced on the NovaSeq 6000 sequencer (Illumina, San Diego, CA) to obtain at least 50,000 reads per spot.

### Bioinformatic analyses

#### Quality Metrics Analysis and initial subgrouping of major cell types

Pre-processed aligned datasets of cells from Cell Ranger were loaded into RStudio to create Seurat objects (Seurat V5.0.1) [18]. Later, a quality metrics analysis was performed to exclude low-quality cells or cells likely undergoing apoptosis. Cells with the highest and lowest RNA features as well as with high percentage of mitochondrial genes (>18%) were excluded from further downstream analyses. Individual datasets were later normalized using SCTransformation [19]. Unsupervised clustering was performed to identify various cell populations and annotated based on expression of conventionally used synovial population markers. For example, *CD45* and *CD14* were used for myeloid cells [20–22], while non-hematopoietic cells were selected based on their negative expression of *CD45*.

#### Integration of non-hematopoietic cells and myeloid cells

To integrate datasets of non-hematopoietic cells from patients across distinct disease conditions (i.e., FAI and hip OA), anchor-based integration method from Seurat was performed. Using *SelectIntegrationFeatures* command, top 3000 variable features were identified. Seurat objects were used as input to find anchors using *FindIntegrationAnchors* and then determined anchors were employed to integrate all datasets using *IntegrateData* function. *RunPCA* analysis was performed for linear dimensionality reduction. The *ElbowPlot* function was used to identify the percentage of variance explained by each principal component and to determine the number of dimensions. Non-linear dimensional reduction of non-hematopoietic cells was performed by Uniform Manifold Approximation and Projection (UMAP) [23]. To further determine heterogeneity of cell composition, *FindNeighbors* and *FindClusters* with resolution of 0.8 were used. We next used *FindConservedMarkers* function to determine conserved non-hematopoietic synovial cell populations between FAI and OA patients. The expression of top 50 genes of each identified conserved cell populations were confirmed using *VlnPlot* and *FeaturePlot* functions. To annotate identified populations, Gene Ontology (GO) functional analysis was performed whereby the top 50 genes from each conserved cell population were uploaded to DAVID bioinformatic database [24, 25]. The same integration pipeline was carried out for myeloid cells from FAI and OA patients. Next, we identified condition-specific unique gene sets for each cell cluster using *FindMarkers* function.

#### Differentiation trajectory and transcription factor binding motif analyses

Pseudotemporal ordering and differentiation trajectory analyses were accomplished using Monocle 3 in RStudio [23, 26–28]. Imported scRNA-seq datasets of non-hematopoietic cells underwent preprocessing and non-linear dimensionality reduction using tSNE. Functions *learn_graph* and *order_cells* were used to construct differentiation trajectories and order cells along pseudotime. Based on previous studies, *DPP4^+^PI16^+^* sublining FLS were used as a starting point of the cell differentiation [22, 29]. Monocle 3 analysis predicted that *EREG^+^CCL20^+^MMP3^hi^* FLS were ordered along the trajectory that originated from *DPP4^+^PI16^+^* cells. Using *find_gene_modules* with pre-set resolution of 0.0002, k value of 7 and max_components of 4, similar gene sets were grouped into different modules. The modules with high and unique gene expression enriched in *EREG^+^CCL20^+^MMP3^hi^* cluster were selected. We next imported the genes from these modules into RcisTarget R package to perform the DNA binding motif analysis [30].

#### Integration of scRNA-seq and Spatial-seq datasets

To determine spatial locations of heterogenous synovial cell populations identified by scRNA-seq, pre-processed Seurat objects of non-hematopoietic scRNA-seq datasets as well as Spatial-seq data of FAI and hip OA synovium samples were normalized using SCTransform, respectively. By using *FindTransferAnchors* and *TransferData* functions, Seurat uses an anchor-based approach to determine integration anchors using populations of non-hematopoietic cells from scRNA-seq datasets as a reference and Spatial-seq datasets as a query set. By setting transferred annotations as a default assay and using *SpatialFeaturePlot* function, we can visualize the spatial location of heterogenous synovial cell populations in the synovium. The similar pipeline was performed to integrate the myeloid cell scRNA-seq datasets with associated Spatial-seq data. Synovial cell populations that were successfully integrated and identified in the Spatial-seq data were further selected for intercellular communication analysis.

#### Intercellular communication analysis

We next used MultiNicheNet R package [31] to investigate how OA progression alters cell-cell interactions between non-hematopoietic and myeloid cells. We first merged associated Seurat objects, and then the matrix of raw counts was used to generate a pseudobulk count matrix for each cell type. Next, analysis of differentially expressed genes (DEGs) was performed on generated matrices using edgeR [31, 32]. Pseudobulk gene expression was later normalized using log-transformation. The set of upregulated and downregulated genes was identified for each receiver cell type across different conditions. MultiNicheNet ligand activity was calculated based on the output from the DEGs. The default logFC thresholds of 0.5 and -0.5, and *p* < 0.05 were set to filter significantly upregulated or downregulated genes for further analysis. For each ligand-receptor-contrast pair, downstream target gene inference was estimated using *get_weighted_ligand_target_links* function. To rank the order of ligand-receptor pairs, the differential expression of ligand and receptor in sender and receiver cells, respectively, and the ligand activity in receiver cells were calculated. The log-normalized pseudobulk expression was derived for ligands and receptors in sender and receiver cells per disease condition to identify the ligands and receptors with high expression levels across all samples. Final prioritization scores were estimated based on calculated weighted aggregation of previously identified parameters including log-normalized pseudobulk expression, differential expression of ligands and receptors, and MultiNicheNet ligand activity. The prioritization scores were used to rank each sender-receiver-ligand-receptor-contrast combination. Finally, across-sample correlation between ligand-receptor pairs and inferred downstream target genes were calculated.

#### Identification of distinct and similar synovial cell populations among hip OA, knee OA, and knee RA patients

To determine if there were conserved and unique synovial cell populations among hip OA, knee OA, and knee RA patients, we applied publicly available knee OA synovium datasets from Chou et al. (GSE152805, n = 3 knee OA samples) [21]. The non-hematopoietic and myeloid cells from knee OA synovial datasets were subsetted, processed, and integrated following our above-mentioned pipelines. For RA synovial cell populations, previously generated heatmaps of myeloid and stromal cell populations from Zhang and co-workers’ study (n = 73 RA and 9 OA samples from different joints) were used with permission from the journal [22]. Note that marker genes that were used to identify synovial cell populations in knee OA and RA datasets were applied to generate heatmaps of myeloid and stromal cell populations in the current hip OA datasets, respectively.

### Immunofluorescent staining and histology

Formalin-fixed paraffin-embedded (FFPE) sections of human FAI and OA synovium were placed in a 60°C oven for three hours and hydrated following standard histological protocol [33]. For antigen retrieval, slides were incubated in the citrate buffer solution (10mM Sodium Citrate, 0.05% Tween® 20 Detergent, #P1379, Sigma-Aldrich, pH 6.0) for 20 min at 95°C. The slides were then incubated in 10% normal goat serum (#S26-100ML, Sigma-Aldrich, diluted with 1X Phosphate-Buffered Saline) for 1 hour at RT. The slides were then incubated in the mixture of primary antibodies against CD16 (100X, #ab201340, Abcam) and COL1A1 (100X, #BS-0578R, Bioss) overnight at 4°C. The following day, slides were thoroughly washed in 1X PBST (1X Phosphate-Buffered Saline, 0.05% Tween® 20 Detergent, #P1379, Sigma-Aldrich) once, and 1X PBS (Phosphate-Buffered Saline) twice. The slides were then incubated in the mixture of Alexa Fluor 555 (100X, #ab150114, Abcam) and Alexa Fluor 488 (250X, #A11008, ThermoFischer) conjugated secondary antibodies for two hours. Slides were thoroughly washed with PBS and sections were incubated with EREG primary antibody (50X, #PA5-119096, ThermoFischer) overnight at 4°C. Note that EREG primary antibody was conjugated with Alexa Fluor 647 using Antibody Labeling Kit (#A20186, ThermoFischer) according to the manufacturer’s instructions. The slides were cover-slipped using VECTASHIELD PLUS Antifade Mounting Medium with DAPI (#H-2000-10, Vector Laboratories) and stored at 4°C. Fluorescent images were taken by an Axioscope 5 (Zeiss) with an Axiocam 305 mono (Zeiss). 8 μm-thick FFPE synovial tissue sections were also stained following standard hematoxylin protocol to visualize cell morphology. Furthermore, the proximity of *EREG^+^* FLS and *CD68^+^COL1A1^+^* fibrotic MΦ was defined if the cell-cell distance between these two populations was ≤ 5 𝜇m apart from each other.

### Statistical Analysis

The Shapiro-Wilk test was used for normality testing. The percentage of positively immunofluorescent stained synovial cells between human FAI and OA patients was compared using unpaired Student t-test. To compare the percentage of myeloid and non-hematopoietic synovial cell populations across different disease conditions and sex, we used two-way ANOVA with Fischer’s LSD test with significance reported at the 95% confidence level. Statistical analyses were performed using GraphPad Prism 10 (GraphPad Software Inc, San Diego, CA). Data were presented as mean±SD. The default statistical analyses embedded in Seurat, Monocle 3, RcisTarget, MultiNicheNet, and GO Term were used for scRNA-seq and Spatial-seq datasets accordingly.

## Results

Hip synovial cells of six Cam-type FAI and six hip OA patients (secondary to FAI) were submitted to UR Genomics Research Center for scRNA-seq analysis (**Supplemental Table 1 and Supplemental Fig. 1**; n = 3 per sex). Unsupervised clustering of scRNA-seq analysis identified 16 and 20 unique cell populations in FAI and hip OA synovium, respectively (**Supplemental Fig. 2)**. These cell populations were further grouped into 6 major cell populations that were conserved both in FAI and hip OA synovium. The 6 conserved major populations were annotated based on their enriched markers and literature [16, 34–36] : 1) *PDPN^+^* FLS, 2) *PECAM1^+^* endothelial cells (ECs), 3) *MCAM^+^* mural cells, 4) *CD45^+^CD14^+^* myeloid cells, 5) *CD45^+^CD3D^+^* T and natural killer (NK) cells, as well as 6) *CD45^+^KIT^+^* mast cells. Next, *CD45^+^CD14^+^* myeloid cells and *CD45^-^* non-hematopoietic cells were examined in downstream analyses.

### Hip OA synovium exhibited increased inflammation and metabolic imbalance compared to FAI synovium

Integrated analysis of *CD45^-^* non-hematopoietic cells (n = 66,658 cells) identified 3 major conserved subpopulations of FLS between FAI and OA synovium: *PRG4^hi^THY1^-^* lining *FLS* (cluster F-0, F-1, and F-12), *PRG4^lo^THY1^+^* sublining *FLS* (clusters F-3, F-4, F-6, and F-8), and *PRG4^med^THY1^-^* transitional *FLS* (cluster F-5 and F-11) (**Fig. 2A, and Supplemental Fig. 3**). We also identified a population of *COL2A1^+^COL1A1^+^* fibrotic chondrocytes (cluster F-2) and a population of *MKI67^+^TOP2A^+^* proliferating cells (cluster F-13). Based on high expression of MCAM (CD146), *ACTA, and RGS5*, we annotated this population as *ACTA^+^RGS5^+^* mural cells (cluster F-10). Interestingly, male patients had significant higher percentage of *ACTA^+^RGS5^+^* mural cells than female OA patients, regardless of disease stage **(Supplemental Fig. 3)**. We also observed three subpopulations of *PECAM1^+^* ECs: *NOTCH4^+^CLND5^+^EDN1^+^*arteriolar ECs (cluster F-7), *DLL4^+^TLL1^+^* ECs (cluster F-9), and *TNF^+^IL1B^+^*pro-inflammatory ECs (cluster F-14) [22]. Hip OA synovium had a higher percentage of *PRG4^+^EREG^+^*FLS versus FAI synovium (**Fig. 2B**, 24% vs. 0.78%; yellow color indicates *PRG4^+^EREG^+^* FLS; green color indicates *PRG4^+^* FLS; orange color indicates *EREG^+^* FLS). Epiregulin (EREG) is one of the ligands of EGFR signaling [37], while lubricin (PRG4) is a common marker for lining FLS. Furthermore, we observed that percentages of *EREG^+^CCL20^+^MMP3^hi^* (cluster F-1) and *CHD17^+^SYT16^+^* lining FLS (cluster F-12) were significantly increased in human hip OA synovium versus those in FAI synovium, regardless of sex (**Fig. 2C**). *PRG4^+^EREG^+^* FLS in OA synovium highly expressed proinflammatory cytokines (*CXCL1* and *CXCL8*) and degrading enzymes (*MMP1* and *MMP3*) compared to the same population in FAI synovium (**Fig. 2D-E**). Conversely, FAI synovium had an increasing trend in cell percentages for *DPP4^+^PI16^+^*, *CD34^+^C3^+^CXCL14^+^*, and *EGR1^+^FOS^+^IER2^+^* sublining FLS as well as fibrotic chondrocytes **(Supplemental Fig. 4)**. Interestingly, male FAI and OA patients had a significantly higher percentage of synovial mural cells than female patients (**Supplemental Fig. 4**). The results of GO functional analysis of each cluster as well as DEGs of the same cell cluster between FAI and OA conditions were listed in Supplemental Files 1 and 2, respectively.

**Fig. 2.**
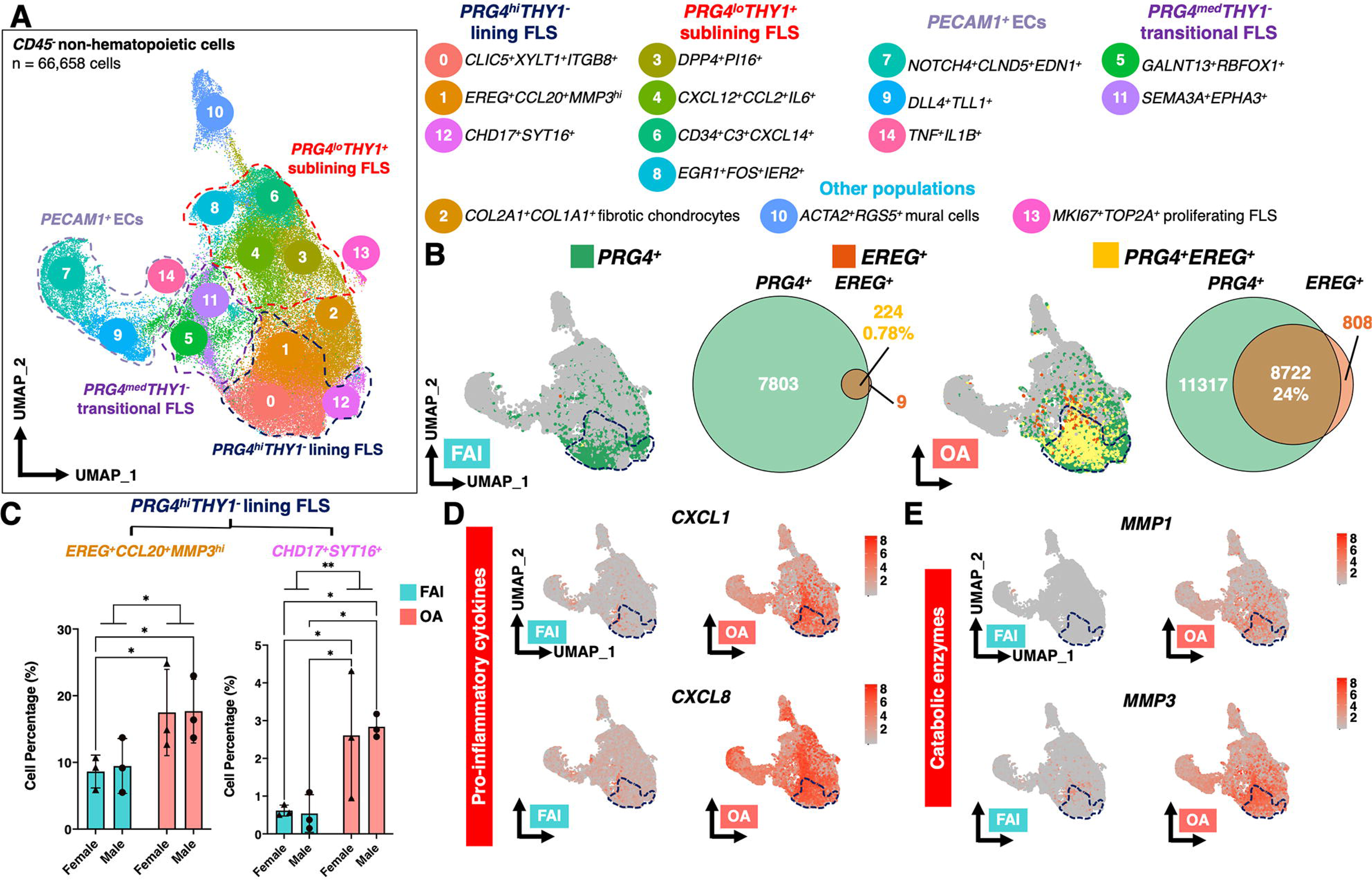
**(A)** Integrated *CD45^-^* non-hematopoietic cells (n = 66,658 cells) from FAI (n = 6; 3 females) and hip OA (n = 6; 3 females) synovium revealed 15 cell subsets that can be grouped into 4 major cell groups: 1) *PRG4^hi^THY1*-lining FLS, 2) *PRG4^lo^THY1^+^* sublining FLS, 3) *PECAMP1^+^* ECs, and 4) *PRG4^med^THY1^-^* transitional FLS. The detailed annotation for each cell subset is also listed. **(B)** Hip OA synovium had significantly increased *PRG4^+^EREG^+^* lining FLS compared to FAI synovium (i.e., yellow-colored cells, 24% in OA vs 0.78% in FAI, normalized to total cell count per disease condition). Interestingly, **(C)** *PRG4^hi^THY1^-^* lining FLS including *EREG^+^CCL20^+^MMP3^hi^* FLS (F-1 cluster) and *CHD17^+^SYT16^+^* FLS (F-12 cluster) were significantly upregulated in OA vs. FAI synovium. Mean±SD. Two-way ANOVA with Fischer’s LSD test, *p <0.05, **p < 0.01. *PRG4^hi^THY1^-^* lining FLS also expressed genes enriched with **(D)** pro-inflammatory cytokines (CXCL1 & CXCL8) and (E) degradation enzymes (MMP1 & MMP3) in OA synovium.

### FAI and hip OA synovial synovium exhibited distinct cellular composition of myeloid cells

We identified 7 major conserved *CD45^+^CD14^+^* myeloid populations (n = 9813 cells) between FAI and hip OA synovium: 1) *IL1B^lo^TNF^hi^* MΦ (clusters M-0 and M-1), 2) *IL1B^hi^IL10^+^* hybrid macrophages (MΦ, clusters M-2, M-3, M-5), 3) *COL1A1^+^* fibrotic MΦ populations (clusters M-8 and M-11), 4) *CD1c^+^* dendritic cells (clusters M-6 and M-9), 5) *MERTK^+^LYVE1^+^* MΦ (cluster M-4), 6) *LTB^+^CD69^+^* myeloid cells (cluster M-7), and 7) *S100A8^+^S100A9^+^* monocytes (cluster M-10) (**Fig. 3A & Supplemental Fig. 5**) [16, 34–36]. Note that MΦ clusters M-0, M-1, M-2, M-3, and M-4 were enriched with CD68 expression (**Supplemental Fig. 5**) The population of *S100A8^+^S100A9^+^* monocytes highly expressed inflammation-triggering alarmins such as *S100A8/9/12*, while *LTB^+^CD69^+^* myeloid cells highly expressed lymphotoxin β (LTB, members of tumor necrosis factor-superfamily) (**Supplemental Fig. 5**). Cell percentages of *S100A8^+^S100A9^+^* monocytes and *LTB^+^CD69^+^* myeloid cells were notably increased in hip OA synovium relative to those in FAI (**Fig. 3B & Supplemental Fig. 6**). Additionally, female OA patients had significant higher percentage of *LTB^+^CD69^+^* myeloid cells than male OA patients. *COL1A1^+^IGFBP5^+^* fibrotic MΦ (cluster M-11) also showed an upward trend in cell percentage in OA synovium. In contrast, *MERTK^+^LYVE1^+^* MΦ (cluster M-4) were increased in FAI synovium, though it was not statistically significant. Furthermore, the results of GO functional analysis of each myeloid cluster as well as DEGs of the same cell cluster between FAI and OA conditions were listed in Supplemental Files 3 and 4, respectively.

**Fig. 3.**
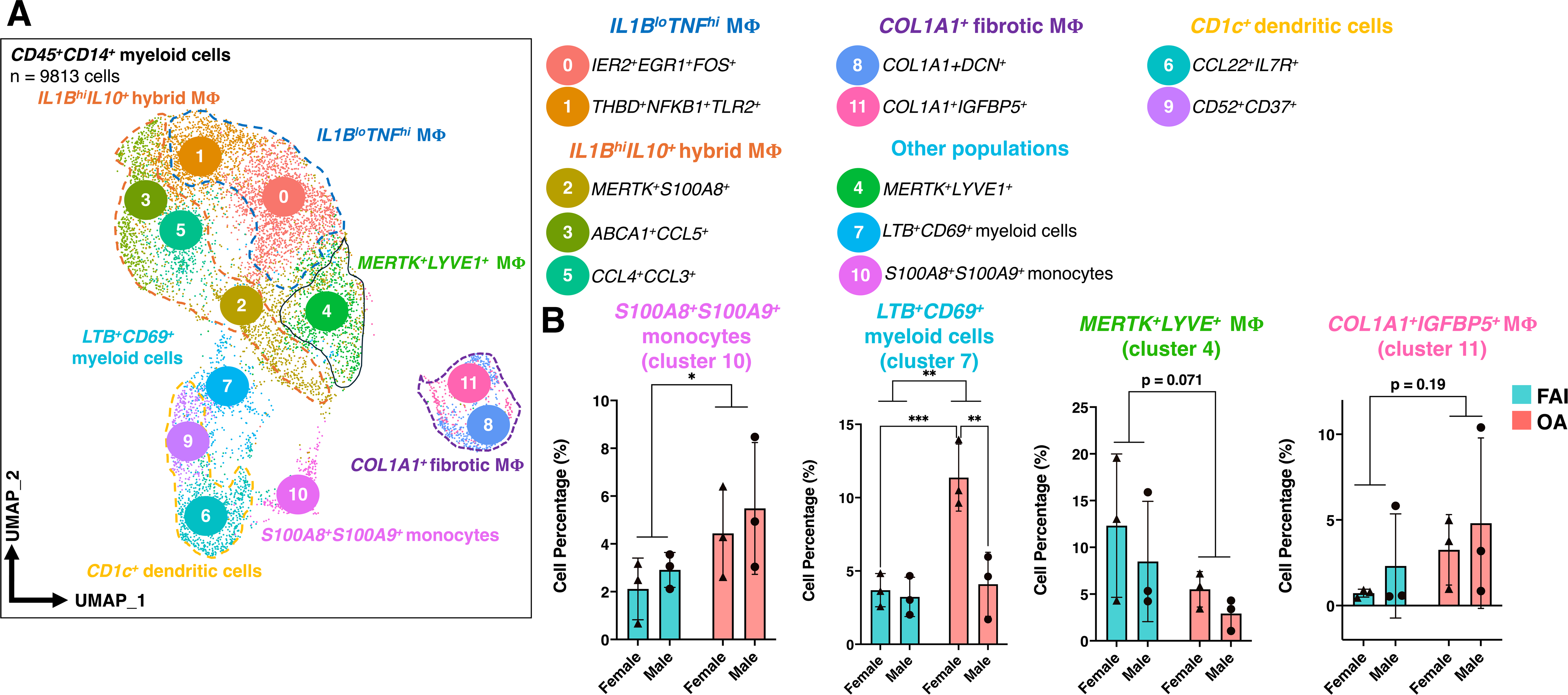
**(A)** Unsupervised clustering of integrated *CD45^+^CD14^+^* myeloid cells (n = 9813 cells) revealed 7 major cell groups in FAI (n = 6, 3 females) and hip OA (n = 6, 3 females) synovium samples: 1) *IL1B^lo^TNF^hi^* MΦ, 2) *COL1A1^+^* fibrotic MΦ, 3) *CD1c*^+^ dendritic cells, 4) *IL1B^hi^IL10^+^* MΦ, 5) *MERTK^+^LYVE1^+^*MΦ, and 6) *LTB^+^CD69^+^*myeloid cells, and 7) *S100A8^+^S100A9^+^* monocytes. **(B)** *LTB^+^CD69^+^*myeloid cells (cluster 7) and *S100A8^+^S100A9^+^*monocytes (cluster 10) were significantly upregulated in hip OA synovium. Hip OA synovium also had a trend toward increased cell percentage of *COL1A1^+^IGFBP5^+^* fibrotic MΦ (cluster 11), while exhibiting a trend toward decreased cell percentage of *MERTK^+^LYVE1^+^* MΦ (cluster 4) (Two-way ANOVA with Fisher LSD’s test, * p < 0.05, ** p < 0.01.)

### *EREG^+^CCL20^+^MMP3^hi^* lining FLS may derive from *DPP4^+^PI16^+^* sublining FLS

We next used Monocle 3 R package to determine the cellular origin of increased pro-inflammatory *PRG4^+^EREG^+^* lining FLS in hip OA synovium [26, 27]. Pseudotime analysis predicted that *DPP4^+^PI16^+^* sublining FLS (cluster F-3) could be a potential source for *EREG^+^CCL20^+^MMP3^hi^* lining FLS [22] (**Fig. 4A**). Further analysis indicated that Pre-B-Cell Leukemia Homeobox 3 (PBX3), Lysine Demethylase 5A (KDM5A), and E2F Transcription Factor 4 (E2F4) might be involved in regulating differentiation of *DPP4^+^PI16^+^* sublining FLS into *EREG^+^CCL20^+^MMP3^hi^* lining FLS (**Fig. 4B**). Expression of *DPP4* and *PI16* decreased, while expression of *PRG4* and *EREG* increased, over differentiation trajectory (**Fig. 4C-D)**. Moreover, PBX3 and KDM5A expression decreased in *EREG^+^CCL20^+^MMP3^hi^* lining FLS during differentiation **(Fig. 4B**). Although the percentages of cells expressing E2F4 increased over the course of differentiation, E2F4 expression levels did not change remarkably between *DPP4^+^PI16^+^* sublining FLS and *EREG^+^CCL20^+^MMP3^hi^* lining FLS **(Fig. 4E**).

**Fig. 4.**
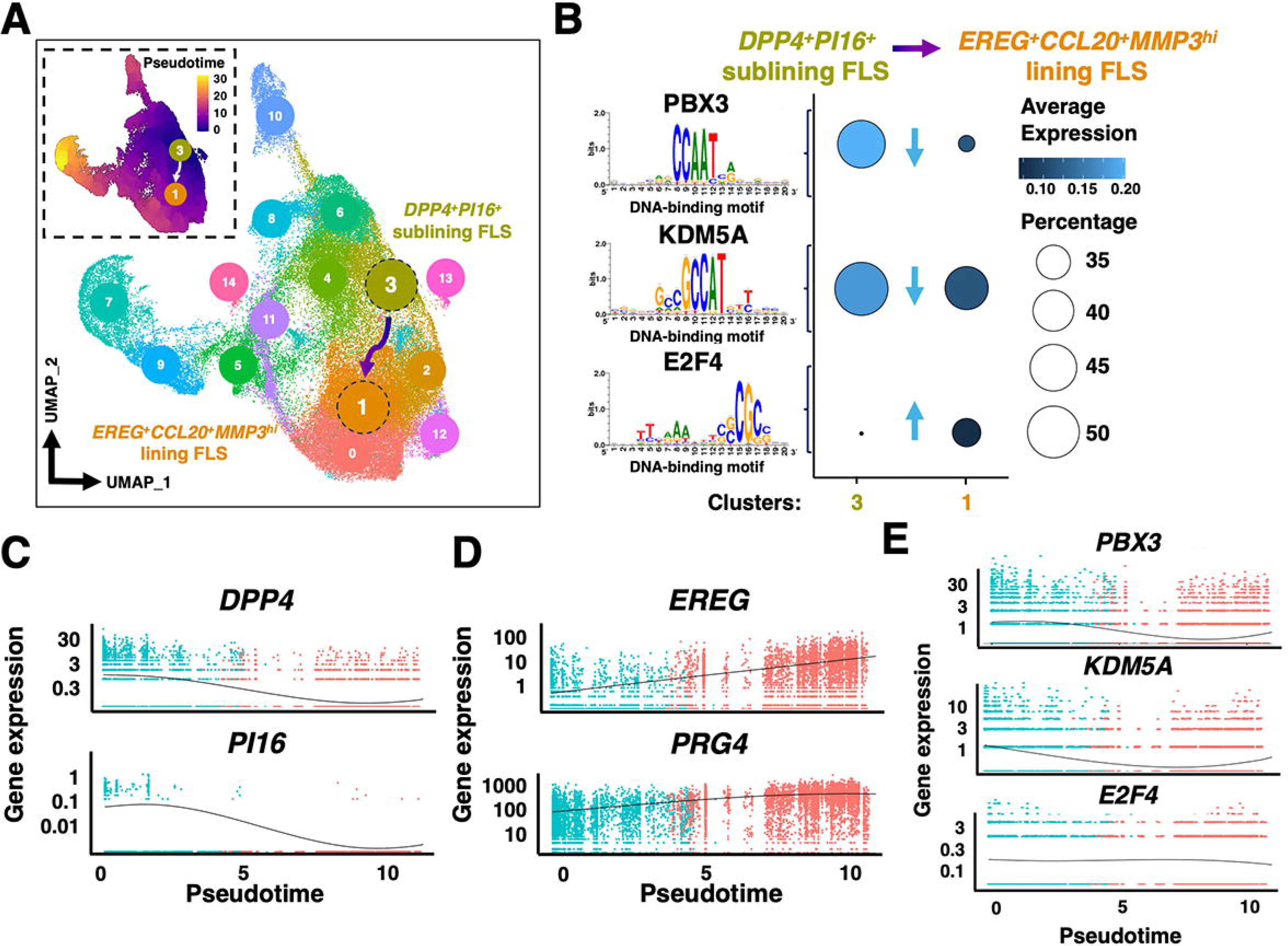
**(A)** Pseudotime analysis with *DPP4^+^PI16^+^* sublining FLS (cluster F-3) predicted by Monocle 3. Arrow indicates differentiation trajectory towards *EREG^+^CCL20^+^MMP3^hi^* lining FLS (cluster F-1). **(B)** DNA-binding motifs analysis identified PBX3, KDM5A, and E2F4 may regulate the differentiation of *DPP4^+^PI16^+^* sublining FLS into *EREG+CCL20^+^MMP3^hi^* lining FLS. In particular, The percentages of cells expressing PBX3 and KDM5A decreases (arrow pointing down), while E2F4 expressing cells increases (arrow pointing up) along with the differentiation. Lighter blue color and larger circle size represent higher relative expression. Pseudotime regression plots for genes associated with **(C)** the *DPP4^+^PI16^+^* perivascular sublining FLS (cluster F-3) (DPP4, PH 6) and **(D)** *EREG+CCL20+MMP3^hi^* lining fibroblasts (cluster F-1) (PRG4, EREG). Furthermore, gene expression of **(E)** Expression levels of PBX3 and KDM5A but not E2F4 changes over pseudotime.

### *COL1A1^+^IGFBP5^+^* fibrotic MΦ may modulate inflammation and cell survival of synovial *EREG^+^CCL20^+^MMP3^hi^* lining FLS through FGF2-SDC4 pathway

To further understand potential intercellular communications, we compared cell-cell crosstalk between lining FLS (clusters F-0, F-1, and F-12) and different MΦ populations (clusters M-1, M-2, M-4, M-7, M-8, M-10, and M-11). These populations were selected because all could be successfully identified across both the scRNA-seq and Spatial-seq datasets. Interestingly, there were strikingly more cell-cell interactions between lining FLS and MΦ in the FAI synovium compared to OA synovium, with *MERTK^+^LYVE1^+^* MΦ (cluster M4) and *LTB^+^CD69^+^* MΦ (cluster M7) serving as the predominant ligand secreting MΦ populations (**Fig. 5A-B**). Conversely, *COL1A1^+^DCN^+^* fibrotic MΦ (cluster M-8) and *COL1A1^+^IGFBP5^+^* fibrotic MΦ (clusters M-11) became major ligand secreting MΦ populations in the OA synovium (**Fig. 5B**). Based on scRNA-seq mapping to Spatial-seq sections, *EREG^+^CCL20^+^MMP3^hi^* lining FLS (cluster F-1) were spatially adjacent to *COL1A1^+^IGFBP5^+^* fibrotic MΦ (cluster M-11) (**Fig. 5C & Supplemental Fig. 7-10**). Although the close vicinity of these two populations was observed in both FAI and hip OA synovium, hip OA patients exhibited remarkably increased co-localization of *EREG^+^CCL20^+^MMP3^hi^* lining FLS and *COL1A1^+^IGFBP5^+^* fibrotic MΦ compared to FAI patients. Moreover, MultiNicheNet analysis revealed that the fibroblast growth factor 2 (FGF2) – syndecan 4 (SDC4) signaling pathway was the most prominent cell-cell interaction between *EREG^+^CCL20^+^MMP3^hi^* lining FLS and *COL1A1^+^IGFBP5^+^* fibrotic MΦ (**Fig. 5B & D**). Furthermore, FGF2-SDC4 signaling was highly upregulated in hip OA synovium (**Fig. 5E)**. The FGF2-SDC4 signaling pathway activated distinct downstream gene sets across different disease states (**Fig. 5E**). GO term analysis of activated genes downstream of FGF2-SDC4 signaling revealed that biological processes associated with aging, inflammation, and angiogenesis were up-regulated in the hip OA synovium, while biological processes involved in mechanical stimuli and skeletal muscle differentiation were dominant in the FAI synovium (**Fig. 5F-G**).

**Fig. 5.**
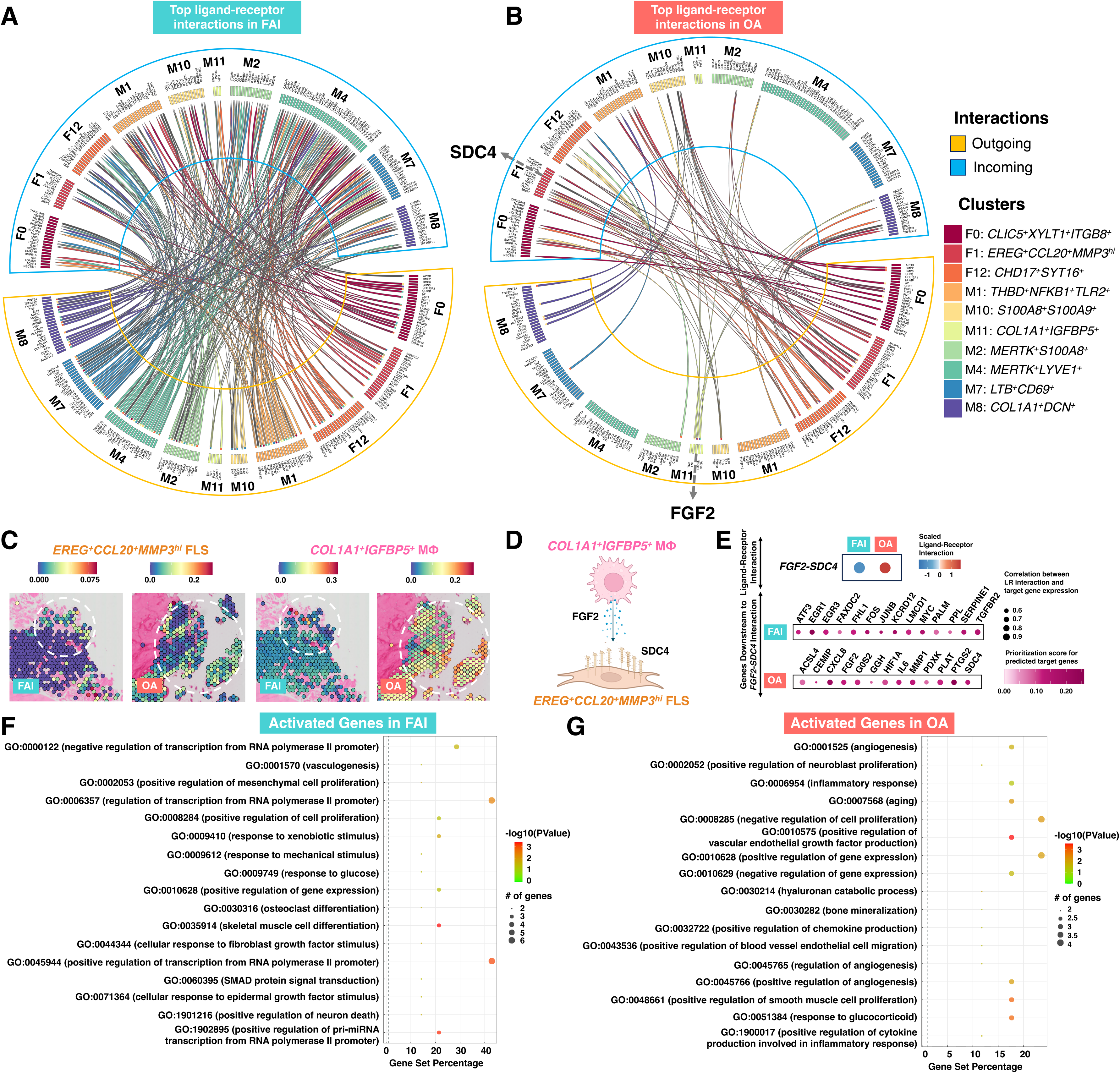
Top ligand-receptor interactions between lining FLS (clusters F-0, F-1, and F-12) and selected MФ (clusters M-1, M-2, M-4, M-7, M-8, M-10, and M-11) in **(A)** FAI and **(B)** hip OA synovium. **(C)** *EREG^+^CCL20^+^MMP3^hi^* lining FLS (F-1) and *COL1A1^+^IGFBP5^+^*fibrotic MФ (M-11) were located close to each other in FAI and OA synovium. Note that there were more *COL1A1^+^IGFBP5^+^* MФ and *EREG^+^CCL20^+^MMP3^hi^* lining FLS in hip OA synovium compared to FAI synovium. **(B, D-E)** *FGF2-SDC4* signaling pathway was the most prominent cell-cell interaction between *EREG^+^CCL20^+^MMP3^hi^*lining FLS and *COL1A1^+^IGFBP5^+^* fibrotic MΦ in hip OA synovium. Predicted activated genes downstream of the signaling pathway between *COL1A1^+^IGFBP5^+^* MФ and *EREG^+^CCL20^+^MMP3^hi^*lining FLS that were upregulated in FAI (**top**) and OA (**bottom**). The darker the magenta color of the dots represents stronger support of activated genes downstream of *FGF2-SDC4* signaling pathway based on prior literature. The larger the size of the dots indicates stronger correlation between Ligand-Receptor interaction and target genes. Gene ontology analysis of downstream activated genes of *FGF2-SDC4* signaling pathway revealed upregulation of distinct biological processes between **(F)** FAI and **(G)** hip OA synovium. Note color and circle size represent p-value and number of genes, respectively.

### The increased proximity of EREG^+^ FLS and CD68^+^COL1A1^+^ fibrotic MΦ, as well as increased EREG^+^ FLS were highly pronounced in lining region of hip OA synovium

Immunofluorescent staining was performed to validate the close vicinity of EREG^+^ FLS (cluster F-1) and CD68^+^COL1A1*^+^* fibrotic MΦ (cluster M-11) (i.e., cell-cell distance is ≤ 5 𝜇m from each other; **Fig. 6A-B and Supplemental Fig. 11-12**). The lining region of hip OA synovium had a significant increase in the cell count of EREG^+^ FLS in comparison to lining region of FAI synovium (**Fig. 6C and Supplemental Fig. 11-12**). However, no significant difference in the percentages of EREG^+^ FLS and CD68^+^COL1A1^+^ fibrotic MΦ in the sublining region of the synovium between FAI and OA were observed. We observed a greater degree of co-localization between EREG^+^ FLS and CD68^+^COL1A1^+^ fibrotic MΦ in the lining area of hip OA synovium and this trend was not observed in sublining region (**Fig. 6A-D**).

**Fig. 6.**
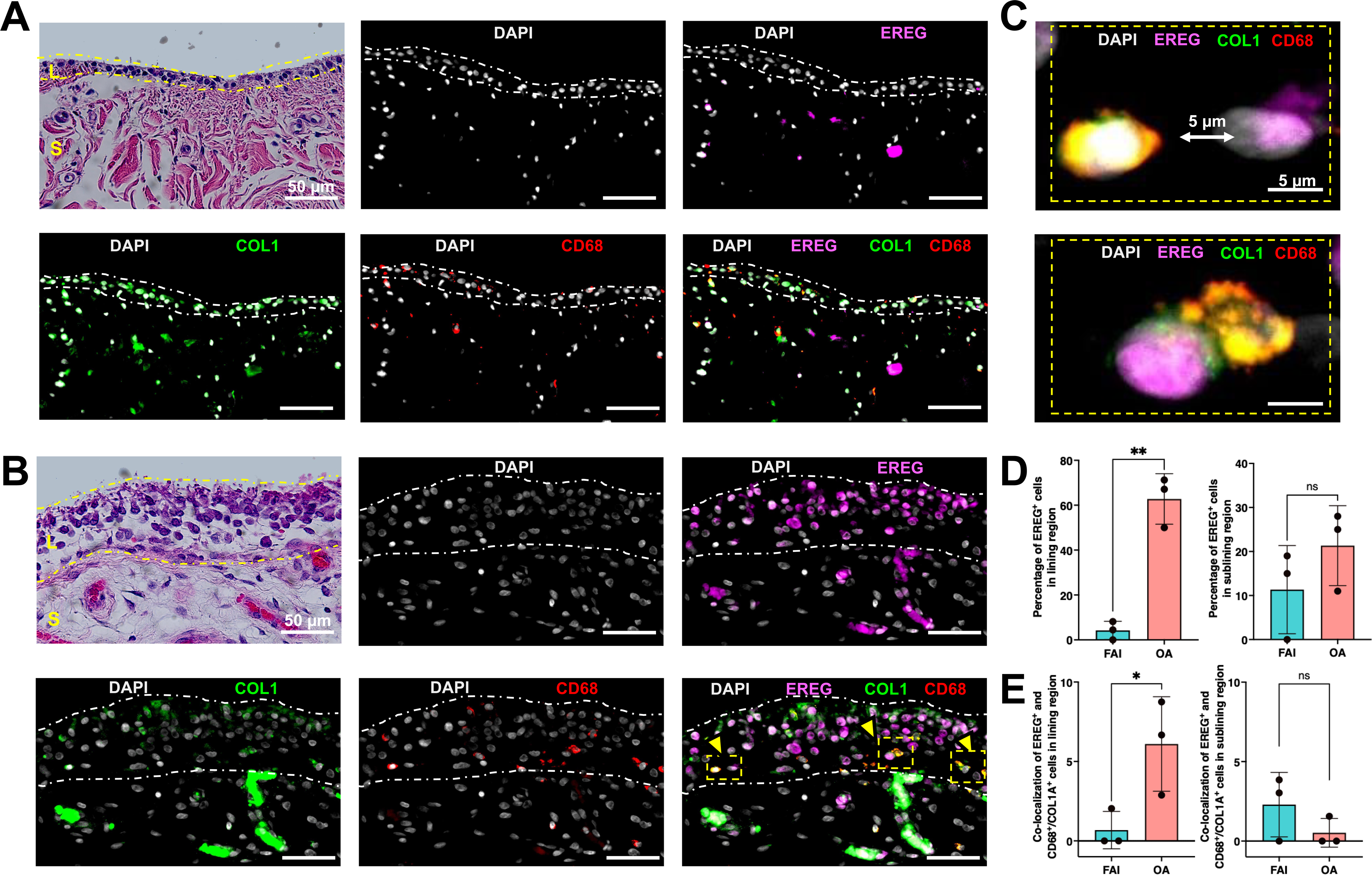
Immunofluorescent staining for EREG, COL1A1, and CD68 in **(A)** FAI and **(B)** hip OA synovium. **L** – lining, **S** – sublining. Yellow arrows indicate the close proximity of EREG+ FLS and CD68^+^ COL1A1^+^ MΦ. Note EREG is expressed mostly by lining FLS in OA synovium. (**C**) High magnification of *EREG*^+^ FLS and CD68^+^ COL1A1^+^ MΦ from (**B**; yellow dashed box). **(D)** Percentage of EREG^+^ FLS within the lining and sublining regions of FAI and OA synovia. **(E)** The numbers of EREG^+^ FLS – CD68+COL1A1^+^ MΦ pairs in lining and sublining regions of FAI and OA synovia. Student’s Unpaired t-test * p < 0.05, ** p < 0.01.

### Distinct and comparable synovial cell populations were detected among hip OA, knee OA, and knee RA patients

Further analysis of DEGs across different clusters revealed novel, previously unreported synovial lining FLS populations that are unique to the synovium of hip OA compared to knee OA and knee RA synovia. For example, *EREG^+^CCL20^+^MMP3^hi^* FLS in the hip OA synovium were uniquely enriched with *IL6*, *CHI3L2*, *CXCL9*, and *IGFBP6* expression, although they also had similar gene expression profiles compared to *PRG4^+^* lining cells and *PRG4^+^CLIC5^+^* lining cells identified by Zhang et al., (knee OA and RA studies) or to intimal FLS identified by Chou et al. (knee OA study) (**Supplemental Fig. 13**) [16, 21, 22]. Conversely, *CXCL12^+^CCL2^+^IL6^+^*, and *CD34^+^C3^+^CXCL14^+^* sublining FLS as well as *ACTA2^+^RGS5^+^* mural cells in hip OA synovium showed similar transcriptomics to *CXCL12^+^SFRP1^+^* and *NOTCH3^+^* sublining FLS or mural cells reported in knee RA synovium [22, 34]. Additionally, the majority of lining FLS identified in our study were enriched with ECM remodeling proteins such as types II, IV, V, and VI collagens, fibronectin, proteoglycans, laminin, and tenascin that are similar to the lining and intimal FLS in knee RA and OA synovium [21, 22, 34] (**Supplemental Fig. 13**). Among hematopoietic cells, *CCL22^+^IL7R^+^* dendritic cells expressed similar gene sets to dendritic cells in knee RA and OA synovia [21, 22], while gene expression of *MERTK^+^S100A8^+^* MΦ (cluster M-2) was comparable to *MERTK^+^S100A8^+^* MΦ in RA synovium [16, 22] (**Supplemental Fig. 14**). Nevertheless, most MΦ and monocytes populations in the hip OA synovium demonstrated unique gene expression that is divergent from synovial MΦ and monocytes in knee OA and RA joints [21, 22] (**Supplemental Fig. 14**).

## Discussion

The findings of this study revealed the dynamic changes in cell composition of synovium between FAI and hip OA patients. Hip synovial FLS can be divided into *PRG4^hi^THY1^-^* lining, *PRG4^lo^THY1^+^* sublining and *PRG4^med^THY1^-^* transitional populations. Sublining FLS populations exhibited high expression of THY1 (CD90), consistent to previous studies showing that THY1 is a common marker for FLS in the sublining region of synovium [20, 22, 34–36]. The GO term analysis indicates that *THY1^+^* sublining FLS and *COL2A1^+^COL1A1^+^* fibrotic chondrocytes (cluster F-2) may be involved in bone formation and articular cartilage development. Interestingly, the *THY1^+^* sublining FLS and fibrotic chondrocytes demonstrates an upward trend in the cell percentage in FAI synovium relative to those in the OA synovium. These findings propound the possibility that sublining FLS and fibrotic chondrocytes likely play a role in the development of the osseous protrusions observed in FAI joints [1, 3–5, 38].

Based on our analyses, EREG is of particular significance in the progression of hip OA. GO term analysis revealed that *EREG^+^CCL20^+^MMP3^hi^* FLS (cluster F-1) are associated with inflammation and angiogenesis, while *CLIC5^+^XYLT^+^* lining FLS (cluster F-0) may be involved in neurogenesis and axon guidance. These findings suggest that *EREG^+^* FLS populations could be related to pain sensitivity in the hip OA joints. Furthermore, our scRNA-seq and immunofluorescent staining results indicate increased *EREG^+^* cells were in the lining region rather than in the sublining region of the OA synovium, implying that there is increased FLS proliferation (i.e., hyperplasia) in the synovial membrane of the hip OA patients. While it is well-recognized that synovial hyperplasia is associated with inflammation in RA [39], our current work further identifies that *EREG^+^* FLS are critical in modulating synovial hyperplasia and inflammation in hip OA. Indeed, elevated expression of *CXCL1* and *CXCL8* as well as cartilage degradation enzymes including *MMP1* and *MMP3* were highly upregulated in *PRG4^hi^EREG^+^* lining FLS of hip OA synovium further suggesting a pro-inflammatory and detrimental role of these cell populations during hip OA development. Only a negligible portion of *PRG4^hi^EREG^+^* lining FLS was observed in FAI synovium. The difference in EREG expression between these disease states indicates that it could be used as a biomarker for hip OA progression. Previous reports show an elevated serum concentration of EREG in RA patients [40]. Additionally, the EREG-mediated temporal expression of growth factors such as amphiregulin, betacellulin, transforming growth factor alpha (TGF-α), and FGF2 has been linked with severe inflammation in arthritis of the knee and ankle joints in mice [40]. Nevertheless, as the etiology is vastly different between OA and RA, the exact molecular mechanisms of EREG-induced inflammation in the hip OA warrant further investigation.

Since our data indicate EREG has great significance in human hip OA development, we next investigated the source of increased *EREG+* cells and their associated signaling pathway. Pseudotime time analysis predicted that *DPP4^+^PI16^+^* sublining FLS may give rise to *EREG^+^CCL20^+^MMP3^hi^* lining FLS via regulation of E2F4, PBX3, and KDM5A. Although the expression level of transcription factor E2F4 is comparable along the pseudotime trajectory, the percentage of cells expressing E2F4 increases as *DPP4^+^PI16^+^* sublining FLS differentiate into *EREG^+^CCL20^+^MMP3^hi^* lining FLS. A recent study reported that E2F2, a member of the E2F transcription factor family similar to E2F4, stimulated CCR4 expression and later induced increased secretion of pro-angiogenic, pro-inflammatory, and degrading factors in synovial FLS of human RA [41]. However, supplementation of PBX3 in C28/I2 human chondrocyte cell line increased the expression of chondrogenic markers such as COL2A1 and ACAN by inhibiting miR-320a overexpression [42]. Finally, despite previous studies reporting that KDM5A is involved in osteoporosis, its role in synovial biology and hip OA is rather limited. Overall, our findings report that *DPP4^+^PI16^+^* cells may serve as a potential cell source for *EREG^+^CCL20^+^MMP3^hi^* lining FLS. This is in line with our previous work demonstrating that *Prg4^hi^* lining FLS arise from a pool of *Dpp4+* progenitors in the synovium of mice with post-traumatic knee OA [29]. Future studies using *in vitro* cell culture and transgenic mouse models are needed to determine the molecular mechanisms by which E2F4, PBX3, and KDM5A regulate differentiation of *EREG^+^CCL20^+^MMP3^hi^* lining FLS in the context of hip OA pathogenesis.

Inflammation is a factor in the development of OA, so we examined the differences in hematopoietic cell populations between FAI and OA samples. Our bioinformatic analyses revealed 12 conserved subtypes of synovial hematopoietic cells in FAI and hip OA patients; however, the percentage of these synovial cells changed remarkably with OA progression. For example, *S100A8^+^S100A9^+^* monocytes with high expression levels of other alarmins, including *S100A9* and *S100A12* were enriched in the OA synovium. It has been shown that S100A8/9/12 secreted by monocytes attract neutrophils and that monocytes induce an increased production of pro-inflammatory cytokines such as TNF and IL6 in resident FLS [16]. Lymphotoxin β (LTB), highly expressed in *LTB^+^CD69^+^* myeloid cells, has been shown to activate follicular dendritic cells, orchestrate the production of T cell-attracting chemokines, and enhance the proliferation of synovial FLS in RA [43]. In FAI synovium, the predominant population *MERTK^+^LYVE1^+^* MΦ (cluster M-4), was previously reported as a tissue resident MΦ population in the sublining layer, and may be involved in perivascular function by supporting collagen production and preventing vascular stiffness [22, 34, 44, 45].

Since sex differences in the development and severity of OA are observed in humans and mice, we paid careful attention to evidence of sexual dimorphism in our datasets. Interestingly, the percentage of hip synovial mural cells was significantly higher in males than in females, irrespective of disease state. This finding highlights a potential sexual dimorphism in the synovial cellular composition during hip OA progression. Although there are few studies regarding sex-dependent alteration in hip synovium, sex-dependent changes in transcriptomic profiles in knee PTOA synovium have been reported. In particular, male mice have been shown to exhibit increased expression of genes involved in inflammation, angiogenesis, fibrosis, and neurogenesis at 28 days following joint injury [46]. Additionally, male knee OA patients exhibited higher synovial vascularization, which is associated with complement activation, in comparison to female patients [47]. Another study reported that synovial fluid from knee OA patients had increased levels of IL2-𝛼, IL3, and TNF-𝛽 in females, while higher levels of MMP1/7/9/13 were observed in the male cohort [48]. Our current work also underpins *ACTA^+^RGS5^+^*mural cells as a potential contributor to sex-specific hip OA development in humans.

This work further identified novel, previously unreported synovial populations (i.e., *CHD17^+^SYT16^+^* lining FLS, *PRG4^med^THY1^-^* transitional FLS, *EGR1^+^FOS^+^IER2^+^* sublining FLS, *MKI67^+^TOP2A^+^* proliferating FLS, and *COL2A1^+^COL1A1^+^* fibrotic chondrocytes) that are unique to hip OA joints. We also found that several synovial cell subsets demonstrate comparable transcriptomic profiles to the synovial cells observed in knee OA and RA patients [16, 17, 20, 22, 34, 35], although whether these cells have the same functionality at different joint sites remains to be elucidated. These findings further underscore the highly heterogeneous nature of synovial cell composition that is not only associated with FAI to OA progression but also varies in a joint-specific manner.

Dysregulation of FGF2 signaling and its downstream gene targets in the FLS may progress hip OA development. Based on our MultiNicheNet analysis, interestingly, a large number of cell-cell crosstalk between lining FLS and different types of MΦ was observed in FAI synovium, while in OA synovium we identified intercellular crosstalk between lining FLS and fibrotic MΦ. Further integration of scRNA-seq and Spatial-seq datasets allowed us to visualize the spatial locations of heterogenous cell populations within the synovium and to investigate the effect of cell-cell crosstalk on hip OA progression. The close vicinity of *EREG^+^CCL20^+^MMP3^hi^* lining FLS and *COL1A1^+^IGFBP5^+^* fibrotic MΦ was highly increased in the hip OA synovium, suggesting potential cell-cell interactions between these two populations. The FGF2/SDC4 signaling pathway was considered as one of the most prominent interactions in hip OA synovium. Our findings suggest that IL6 and IL8 (CXCL8) downstream of this signaling pathway are activated and may be key players involved in synovial inflammation, including enhanced angiogenesis in hip OA synovium (vs. FAI synovium). This aligns with previous studies demonstrating that elevated levels of IL6 and IL8 in synovial fluid have a strong correlation with the radiographic progression of knee OA [49]. Furthermore, elevated levels of FGF2 in synovial fluid are highly correlated with bone destruction and can lead to synovial hyperplasia and cartilage degeneration in human knee RA joints [50]. Despite this, the role of FGF2 in OA development remains controversial. On the one hand, *Fgf2* global knock-out (gKO) mice develop more severe spontaneous and injury-induced OA with elevated expression of a disintegrin and metalloproteinase with thrombospondin motifs 5 (ADAMTS-5) compared to wild-type mice [51, 52]. Contrarily, FGF2 can induce catabolic effects on human cartilage *in vitro* and may disrupt the physiological balance of the production of type I and type II collagens and inhibit the formation of proteoglycans [53, 54]. Interestingly, a recent study also reported that FGF2 may induce expression of ADAMTS5 through p65/miR-105 signaling in human knee OA chondrocytes *in vitro* [53]. The expression of SDC4 by FLS is elevated in RA synovial tissues versus healthy tissues, and silencing of SDC4 via lentiviral short hairpin RNA decreased synovial inflammation in RA [55]. Furthermore, levels of SDC4 in synovial fluid are positively associated with disease progression in knee OA patients and may promote OA severity by increasing the expression of ADAMTS5 [51, 56]. Conversely, the loss of SDC4 in mice precluded the development of OA-like phenotype [51, 57].

A relatively small sample size is a potential limitation of the current study. Nevertheless, our work unveiled critical contributions of synovial cells to hip OA progression from FAI. Future studies may wish to include healthy synovial tissues to construct a comprehensive roadmap of hip OA development in humans. Additionally, investigations using human primary FAI and hip OA synovial cells and/or transgenic mouse models to further validate the findings of the present study are also warranted.

Our work not only elucidated significant changes in the synovial cell populations between FAI and hip OA patients but, most importantly, uncovered the spatial locations and interactions of these cell subsets in the synovium that orchestrate synovial hyperplasia, inflammation, and OA progression. Furthermore, our findings suggest that targeting FGF2-SDC4 signaling between *PRG4^+^EREG^+^* lining FLS and fibrotic MΦ may provide new therapeutic applications for hip OA patients.

## Acknowledgments.

The authors thank UR Genomics Research Center for assisting with scRNA-seq and Spatial-seq. This study was supported in part by NIH grants AR075899, AR082403, UR and C-COMP P30 pilots, Orthopaedic Research and Education Foundation, and Arthritis National Research Foundation.

## References

1. Zhang, C., et al., Femoroacetabular impingement and osteoarthritis of the hip. Can Fam Physician, 2015. 61(12): p. 1055–60.

2. Pun, S., D. Kumar, and N.E. Lane, Femoroacetabular impingement. Arthritis Rheumatol, 2015. 67(1): p. 17–27.

3. Haneda, M., et al., Distinct Pattern of Inflammation of Articular Cartilage and the Synovium in Early and Late Hip Femoroacetabular Impingement. Am J Sports Med, 2020. 48(10): p. 2481–2488.

4. Agricola, R., et al., Cam impingement causes osteoarthritis of the hip: a nationwide prospective cohort study (CHECK). Ann Rheum Dis, 2013. 72(6): p. 918–23.

5. Nasser, R. and B. Domb, Hip arthroscopy for femoroacetabular impingement. EFORT Open Rev, 2018. 3(4): p. 121–129.

6. Agricola, R., et al., Femoroacetabular impingement syndrome in middle-aged individuals is strongly associated with the development of hip osteoarthritis within 10-year follow-up: a prospective cohort study (CHECK). Br J Sports Med, 2024. 58(18): p. 1061–1067.

7. Kamenaga, T., et al., A Novel Model of Hip Femoroacetabular Impingement in Immature Rabbits Reproduces the Distinctive Head-Neck Cam Deformity. Am J Sports Med, 2022. 50(7): p. 1919–1927.

8. Kuhns, B.D., et al., Whole-genome RNA sequencing identifies distinct transcriptomic profiles in impingement cartilage between patients with femoroacetabular impingement and hip osteoarthritis. J Orthop Res, 2023. 41(7): p. 1517–1530.

9. Sanchez-Lopez, E., et al., Synovial inflammation in osteoarthritis progression. Nat Rev Rheumatol, 2022. 18(5): p. 258–275.

10. Sellam, J. and F. Berenbaum, The role of synovitis in pathophysiology and clinical symptoms of osteoarthritis. Nat Rev Rheumatol, 2010. 6(11): p. 625–35.

11. Yuan, J., et al., Genetically predicted obesity and risk of hip osteoarthritis. Eat Weight Disord, 2023. 28(1): p. 11.

12. Murphy, L.B., et al., One in four people may develop symptomatic hip osteoarthritis in his or her lifetime. Osteoarthritis Cartilage, 2010. 18(11): p. 1372–9.

13. Wu, C.L., et al., Dietary fatty acid content regulates wound repair and the pathogenesis of osteoarthritis following joint injury. Ann Rheum Dis, 2015. 74(11): p. 2076–83.

14. Wu, C.L., et al., Serum and synovial fluid lipidomic profiles predict obesity-associated osteoarthritis, synovitis, and wound repair. Sci Rep, 2017. 7: p. 44315.

15. Grieshaber-Bouyer, R., et al., Divergent Mononuclear Cell Participation and Cytokine Release Profiles Define Hip and Knee Osteoarthritis. J Clin Med, 2019. 8(10).

16. Alivernini, S., et al., Distinct synovial tissue macrophage subsets regulate inflammation and remission in rheumatoid arthritis. Nat Med, 2020. 26(8): p. 1295–1306.

17. Zhang, F., et al., Defining inflammatory cell states in rheumatoid arthritis joint synovial tissues by integrating single-cell transcriptomics and mass cytometry. Nat Immunol, 2019. 20(7): p. 928–942.

18. Hao, Y., et al., Dictionary learning for integrative, multimodal and scalable single-cell analysis. Nat Biotechnol, 2024. 42(2): p. 293–304.

19. Hafemeister, C. and R. Satija, Normalization and variance stabilization of single-cell RNA-seq data using regularized negative binomial regression. Genome Biol, 2019. 20(1): p. 296.

20. Mizoguchi, F., et al., Functionally distinct disease-associated fibroblast subsets in rheumatoid arthritis. Nat Commun, 2018. 9(1): p. 789.

21. Chou, C.H., et al., Synovial cell cross-talk with cartilage plays a major role in the pathogenesis of osteoarthritis. Sci Rep, 2020. 10(1): p. 10868.

22. Zhang, F., et al., Deconstruction of rheumatoid arthritis synovium defines inflammatory subtypes. Nature, 2023. 623(7987): p. 616–624.

23. McInnes, L., J. Healy, and J. Melville, Umap: Uniform manifold approximation and projection for dimension reduction. arXiv preprint arXiv:1802.03426, 2018.

24. Huang da, W., B.T. Sherman, and R.A. Lempicki, Systematic and integrative analysis of large gene lists using DAVID bioinformatics resources. Nat Protoc, 2009. 4(1): p. 44–57.

25. Sherman, B.T., et al., DAVID: a web server for functional enrichment analysis and functional annotation of gene lists (2021 update). Nucleic Acids Res, 2022. 50(W1): p. W216–w221.

26. Cao, J., et al., The single-cell transcriptional landscape of mammalian organogenesis. Nature, 2019. 566(7745): p. 496–502.

27. Qiu, X., et al., Reversed graph embedding resolves complex single-cell trajectories. Nat Methods, 2017. 14(10): p. 979–982.

28. Trapnell, C., et al., The dynamics and regulators of cell fate decisions are revealed by pseudotemporal ordering of single cells. Nat Biotechnol, 2014. 32(4): p. 381–386.

29. Knights, A.J., et al., Synovial fibroblasts assume distinct functional identities and secrete R-spondin 2 in osteoarthritis. Ann Rheum Dis, 2023. 82(2): p. 272–282.

30. Aibar, S., et al., SCENIC: single-cell regulatory network inference and clustering. Nat Methods, 2017. 14(11): p. 1083–1086.

31. Browaeys, R., et al., MultiNicheNet: a flexible framework for differential cell-cell communication analysis from multi-sample multi-condition single-cell transcriptomics data. bioRxiv, 2023: p. 2023.06.13.544751.

32. Robinson, M.D., D.J. McCarthy, and G.K. Smyth, edgeR: a Bioconductor package for differential expression analysis of digital gene expression data. Bioinformatics, 2010. 26(1): p. 139–40.

33. Kenney, H.M., et al., Multi-omics analysis identifies IgG2b class-switching with ALCAM-CD6 co-stimulation in joint-draining lymph nodes during advanced inflammatory-erosive arthritis. Front Immunol, 2023. 14: p. 1237498.

34. Kemble, S. and A.P. Croft, Critical Role of Synovial Tissue-Resident Macrophage and Fibroblast Subsets in the Persistence of Joint Inflammation. Front Immunol, 2021. 12: p. 715894.

35. Smith, M.H., et al., Heterogeneity of Inflammation-associated Synovial Fibroblasts in Rheumatoid Arthritis and Its Drivers. bioRxiv, 2022: p. 2022.02.28.482131.

36. Wei, K., et al., Notch signalling drives synovial fibroblast identity and arthritis pathology. Nature, 2020. 582(7811): p. 259–264.

37. Ma, S., et al., Epiregulin confers EGFR-TKI resistance via EGFR/ErbB2 heterodimer in non-small cell lung cancer. Oncogene, 2021. 40(14): p. 2596–2609.

38. Agricola, R., et al., Pincer deformity does not lead to osteoarthritis of the hip whereas acetabular dysplasia does: acetabular coverage and development of osteoarthritis in a nationwide prospective cohort study (CHECK). Osteoarthritis Cartilage, 2013. 21(10): p. 1514–21.

39. Neumann, E., et al., Rheumatoid arthritis progression mediated by activated synovial fibroblasts. Trends Mol Med, 2010. 16(10): p. 458–68.

40. Harada, M., et al., Temporal expression of growth factors triggered by epiregulin regulates inflammation development. J Immunol, 2015. 194(3): p. 1039–46.

41. Xu, W., S. Li, and X. Chang, E2F2 stimulates CCR4 expression and activates synovial fibroblast-like cells in rheumatoid arthritis. Cent Eur J Immunol, 2021. 46(1): p. 27–37.

42. Jin, Y., et al., The role of miR-320a and IL-1β in human chondrocyte degradation. Bone Joint Res, 2017. 6(4): p. 196–203.

43. Tran, C.N., S.K. Lundy, and D.A. Fox, Synovial biology and T cells in rheumatoid arthritis. Pathophysiology, 2005. 12(3): p. 183–9.

44. Knights, A.J., et al., Synovial macrophage diversity and activation of M-CSF signaling in post-traumatic osteoarthritis. bioRxiv, 2023.

45. Lim, H.Y., et al., Hyaluronan Receptor LYVE-1-Expressing Macrophages Maintain Arterial Tone through Hyaluronan-Mediated Regulation of Smooth Muscle Cell Collagen. Immunity, 2018. 49(2): p. 326–341.e7.

46. Bergman, R.F., et al., Sexual dimorphism of the synovial transcriptome underpins greater PTOA disease severity in male mice following joint injury. Osteoarthritis Cartilage, 2023.

47. Sodhi, E.U., et al., Sex-Differences and Associations Between Complement Activation and Synovial Vascularization in Patients with Late-Stage Knee Osteoarthritis. Front Immunol, 2022. 13: p. 890094.

48. Pan, Q., et al., Characterization of osteoarthritic human knees indicates potential sex differences. Biol Sex Differ, 2016. 7: p. 27.

49. Monibi, F., et al., Identification of Synovial Fluid Biomarkers for Knee Osteoarthritis and Correlation with Radiographic Assessment. J Knee Surg, 2016. 29(3): p. 242–7.

50. Manabe, N., et al., Involvement of fibroblast growth factor-2 in joint destruction of rheumatoid arthritis patients. Rheumatology (Oxford), 1999. 38(8): p. 714–20.

51. Pap, T. and A. Korb-Pap, Cartilage damage in osteoarthritis and rheumatoid arthritis--two unequal siblings. Nat Rev Rheumatol, 2015. 11(10): p. 606–15.

52. Chia, S.L., et al., Fibroblast growth factor 2 is an intrinsic chondroprotective agent that suppresses ADAMTS-5 and delays cartilage degradation in murine osteoarthritis. Arthritis Rheum, 2009. 60(7): p. 2019–27.

53. Ji, Q., et al., miR-105/Runx2 axis mediates FGF2-induced ADAMTS expression in osteoarthritis cartilage. J Mol Med (Berl), 2016. 94(6): p. 681–94.

54. Nummenmaa, E., et al., Effects of FGF-2 and FGF receptor antagonists on MMP enzymes, aggrecan, and type II collagen in primary human OA chondrocytes. Scand J Rheumatol, 2015. 44(4): p. 321–30.

55. Cai, P., et al., Syndecan-4 involves in the pathogenesis of rheumatoid arthritis by regulating the inflammatory response and apoptosis of fibroblast-like synoviocytes. J Cell Physiol, 2020. 235(2): p. 1746–1758.

56. Bollmann, M., et al., MMP-9 mediated Syndecan-4 shedding correlates with osteoarthritis severity. Osteoarthritis Cartilage, 2021. 29(2): p. 280–289.

57. Echtermeyer, F., et al., Syndecan-4 regulates ADAMTS-5 activation and cartilage breakdown in osteoarthritis. Nat Med, 2009. 15(9): p. 1072–6.

