## Supplemental Figures for "Integrative scRNA-seq and spatial transcriptomics uncovers distinct macrophage-fibroblast cross-talk in human hip synovium between patients with femoroacetabular impingement and osteoarthritis"

| ID | Disease | Sex | Age (years) | BMI ($kg/m^2$) | Race/ Ethnicity | Experiment |
|---|---|---|---|---|---|---|
| FAI001 | FAI | M | 23 | 22.0 | White | scRNA-seq, Spatial-seq, IF |
| FAI002 | FAI | M | 21 | 27.9 | White | scRNA-seq, Spatial-seq |
| FAI003 | FAI | F | 33 | 21.6 | White | scRNA-seq |
| FAI004 | FAI | F | 18 | 28.1 | White | scRNA-seq, IF |
| FAI005 | FAI | M | 28 | 23.1 | White | Spatial-seq, IF |
| FAI006 | FAI | F | 33 | 28.5 | White | Spatial-seq |
| FAI007 | FAI | M | 32 | 22.7 | White | scRNA-seq |
| FAI008 | FAI | F | 42 | 28.0 | White | scRNA-seq |
| OA001 | OA | M | 66 | 31.3 | White | scRNA-seq, Spatial-seq |
| OA002 | OA | F | 68 | 29.2 | White | Spatial-seq |
| OA003 | OA | F | 62 | 32.4 | White | Spatial-seq |
| OA004 | OA | F | 69 | 31.9 | African American | Spatial-seq |
| OA005 | OA | M | 62 | 28.0 | White | scRNA-seq, Spatial-seq, IF |
| OA006 | OA | M | 60 | 31.0 | White | Spatial-seq, IF |
| OA007 | OA | F | 59 | 39.5 | White | scRNA-seq, IF |
| OA008 | OA | F | 63 | 41.2 | African American | scRNA-seq |
| OA009 | OA | F | 60 | 29.8 | White | scRNA-seq |
| OA010 | OA | M | 66 | 30.0 | White | scRNA-seq |

**Supplemental Table 1 | Patients Demographics.** Synovial biopsies were harvested from the paralabral region of hip joints of Cam-type FAI and hip OA patients (secondary to FAI) undergoing hip arthroscopy and total hip arthroplasty, respectively. **F** – Female, **M** – Male; **IF** – Immunofluorescent staining.

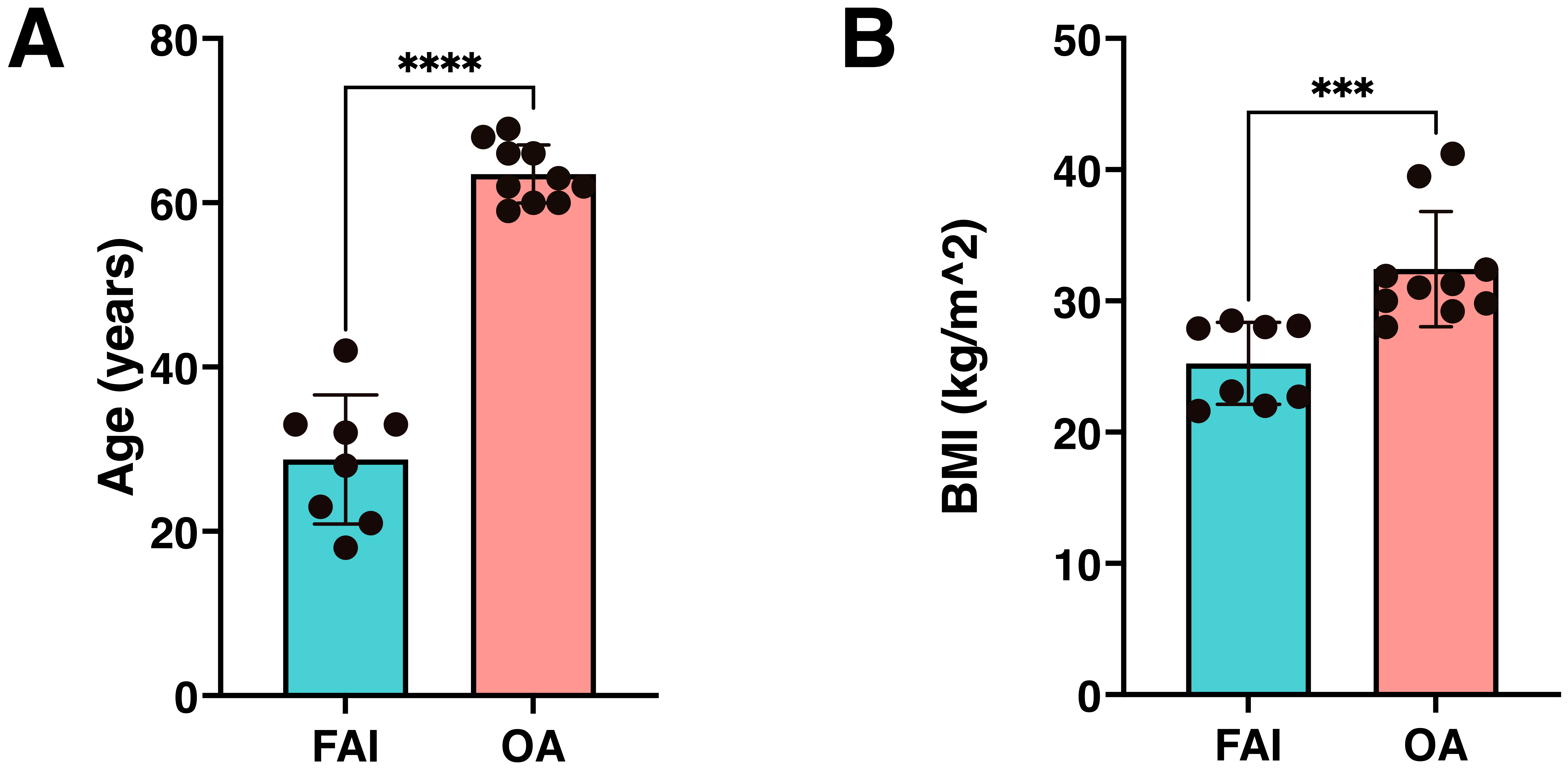


**Supplemental Fig. 1** | Hip OA patient were significantly **(A)** older and had significantly higher **(B)** BMI than FAI patients (Student's Unpaired t-test with Welch's correction. *** $p < 0.001$, **** $p < 0.0001$.)

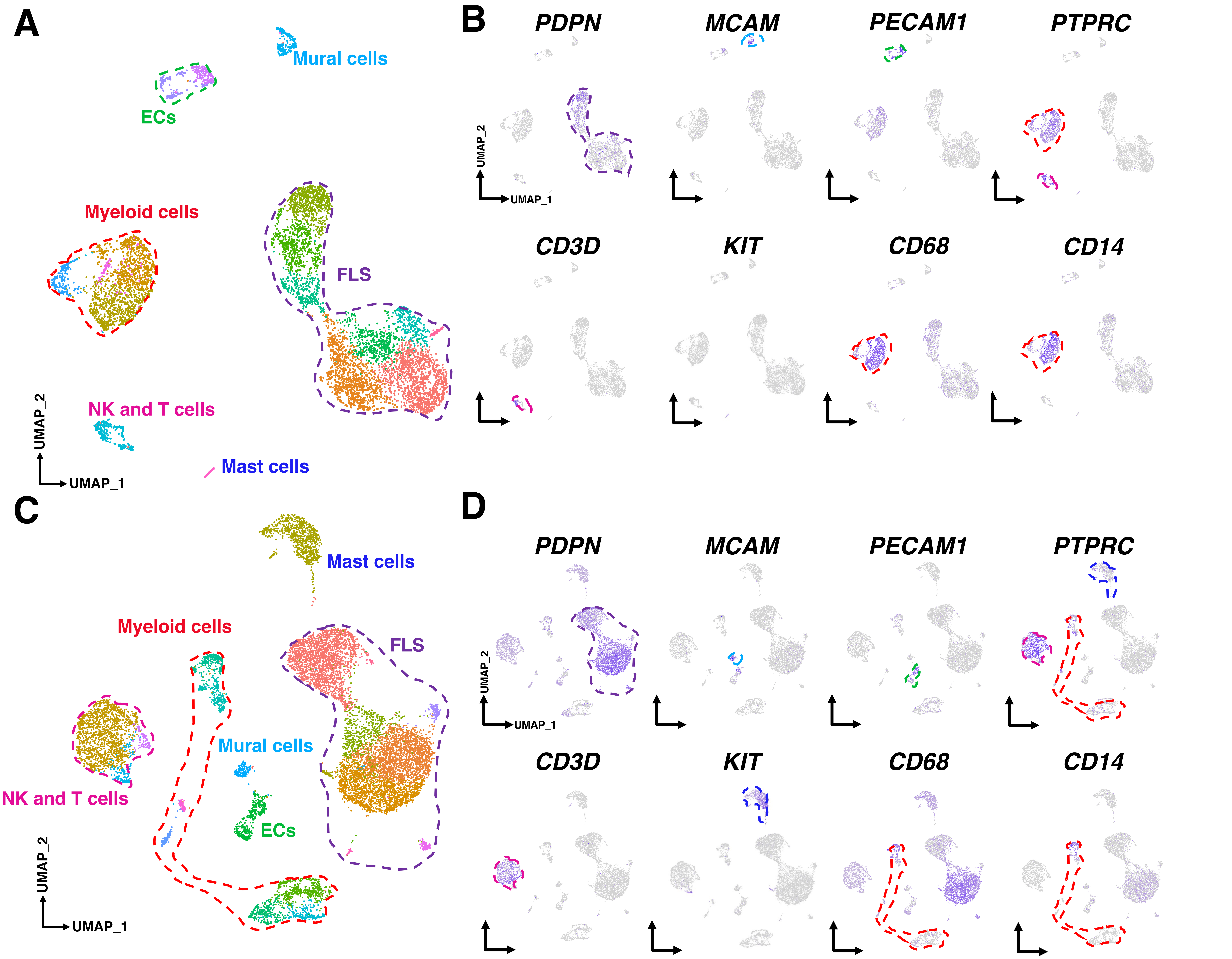


**Supplemental Fig. 2 l Initial unsupervised clustering of scRNA-seq datasets .** (**A**) 16 unique populations in FAI synovium. (**B**) Markers of FAI synovial cell populations. (**C**) 20 unique populations in hip OA synovium**.** (**D**) Markers of OA synovial cell populations. Based on conventional synovial cell markers, 6 conserved major cell populations both in FAI and OA synovium were identified: 1) *CD45⁻PDPN⁺* FLS, 2) *CD45⁻PECAM1⁺* ECs, 3) *CD45⁻MCAM⁺* mural cells, 4) *CD45⁺CD14⁺* myeloid cells, 5) *CD45⁺CD3D⁺* NK and T cells as well as 6) *CD45⁺KIT⁺* mast cells.

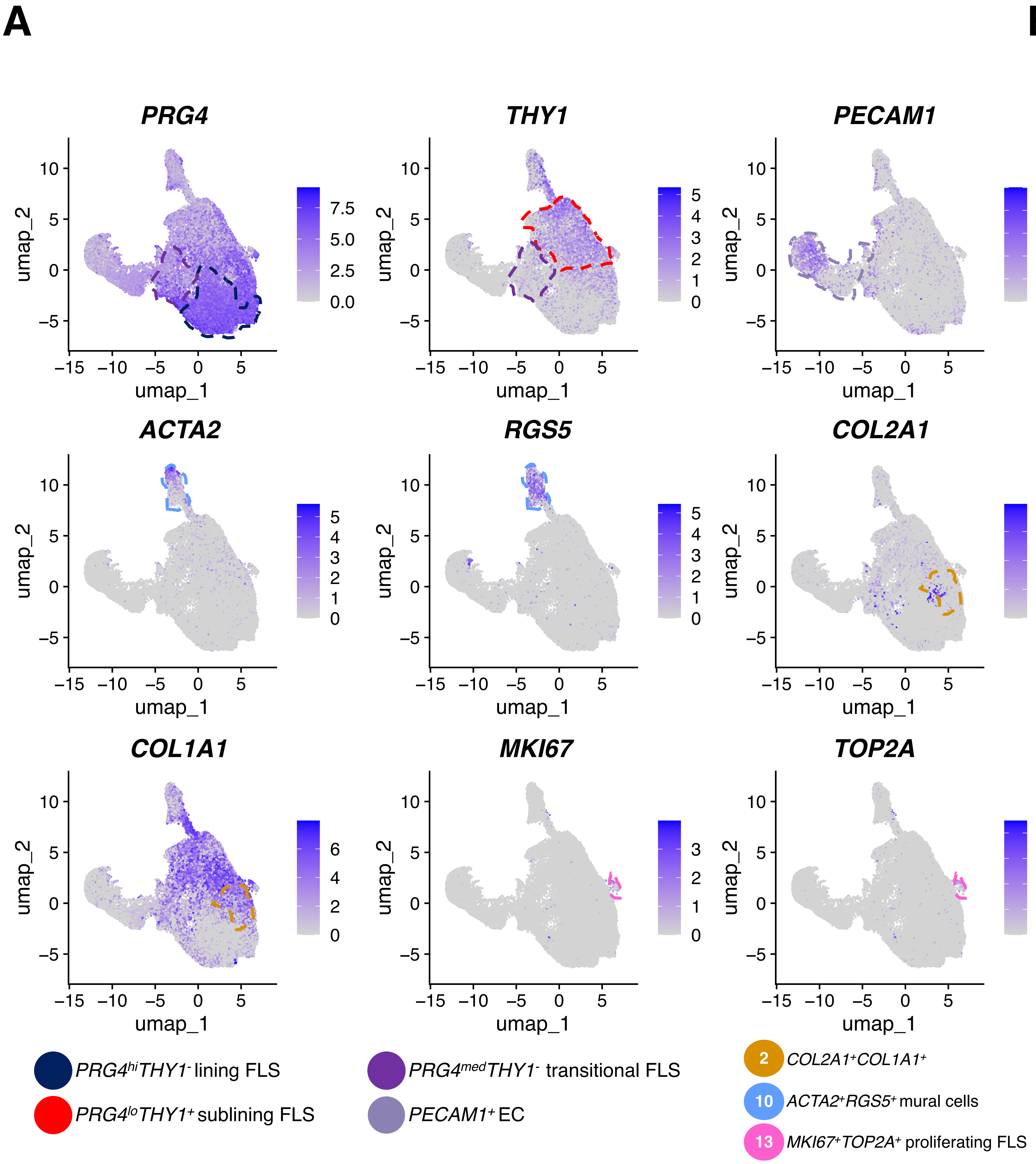


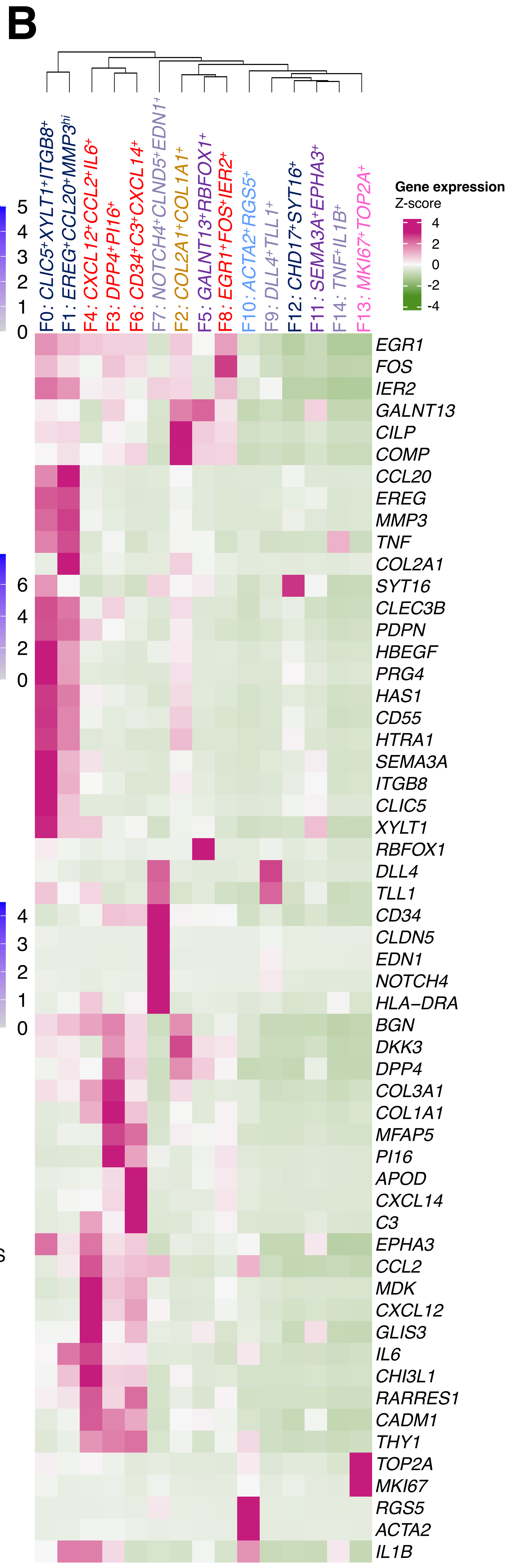


**Supplemental Fig. 3 | Detailed analysis of *$CD45^{-}$* non-hematopoietic cells. (A)** Expression of general markers separating non-hematopoietic cells into *$PRG4^{hi}THY1^{-}$* lining FLS (dark blue dashed line), *$PRG4^{lo}THY1^{+}$* sublining FLS (red dashed line), *$PRG4^{med}THY1^{-}$* transitional FLS (purple dashed line), *$PECAM1^{+}$* ECs (lavender dashed line), *$COL2A1^{+}COL1A1^{+}$* fibrotic chondrocytes (brown dashed line), *$ACTA2^{+}RGS5^{+}$* mural cells (light blue dashed line), and *$MKI67^{+}TOP2A^{+}$* proliferating FLS (pink dashed line). **(B)** Heatmap of finer-grained clusters of non-hematopoietic cells grouped by similarities in normalized expression of differentially expressed genes.

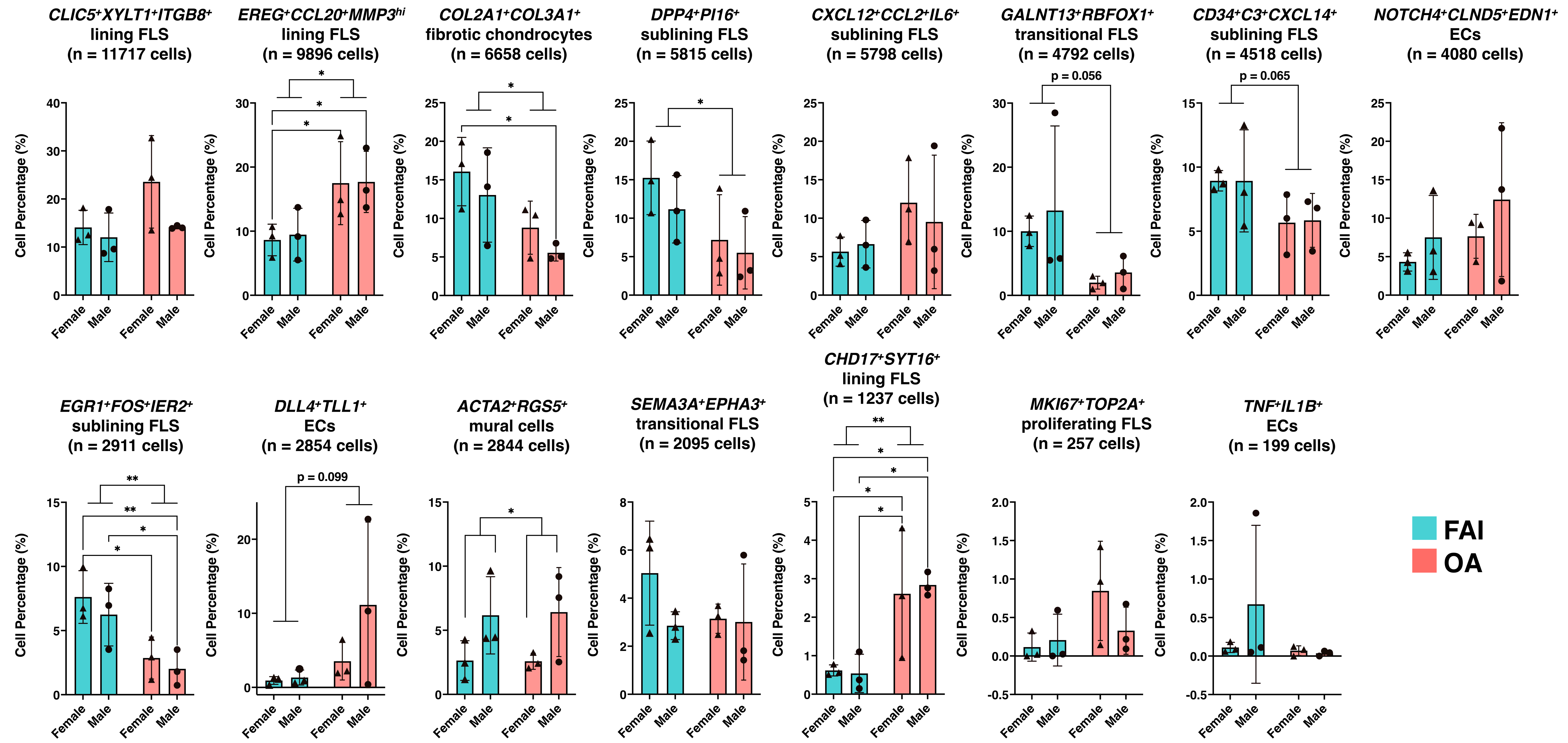


**Supplemental Fig. 4 | Altered percentages of *CD45*⁻ non-hematopoietic synovial cell populations between FAI and OA patients.** Mean±SD. Two-way ANOVA with Fischer's LSD test, *$p < 0.05$, **$p < 0.01$.

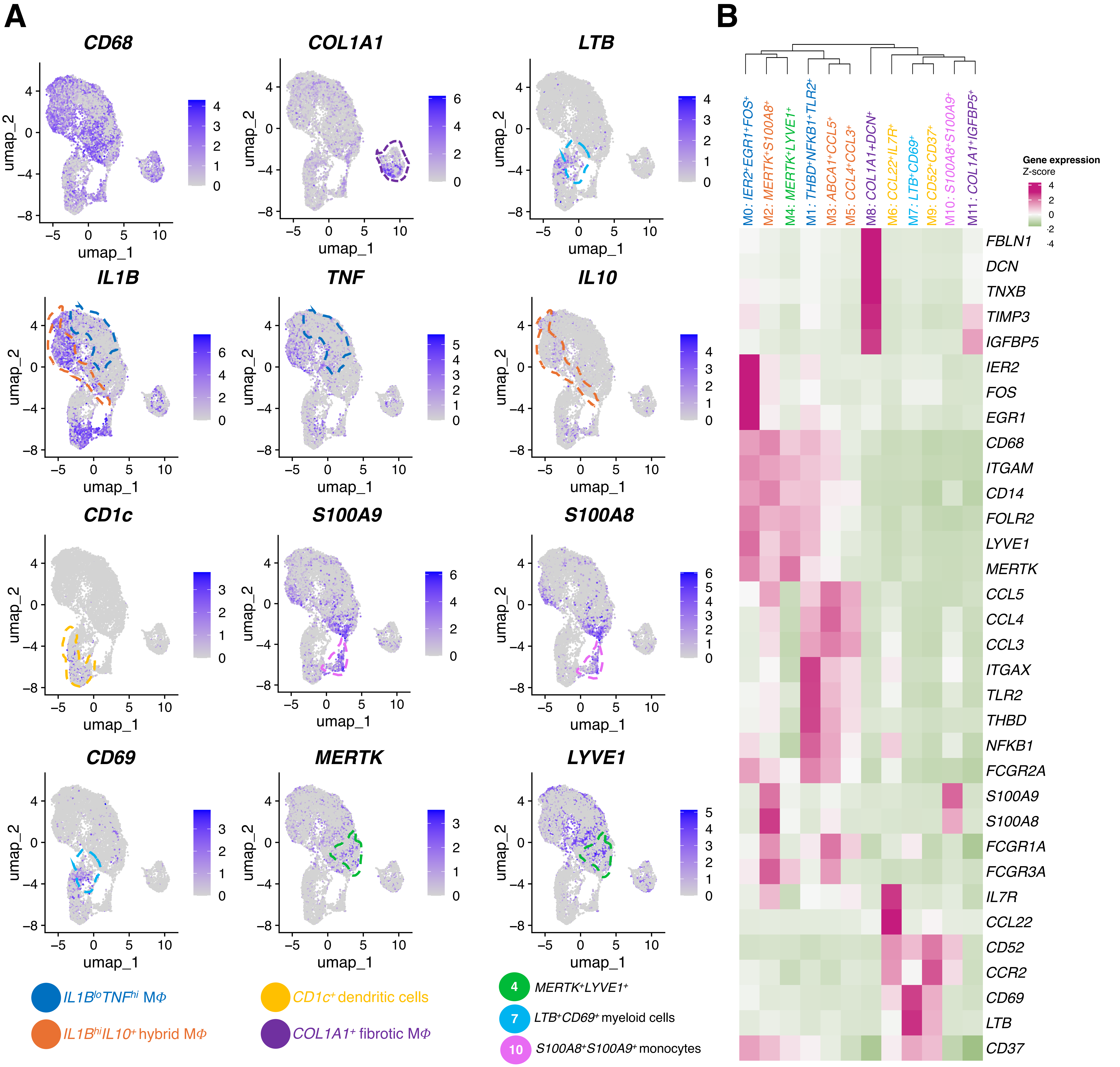


**Supplemental Fig. 5 | Marker gene expression across unique *CD45⁺CD14⁺* myeloid cell subsets. (A)** Expression of marker genes for *IL1B^lo TNF^hi* MΦ (blue dashed line), *IL1B^hi IL10^+* hybrid MΦ (orange dashed line), *CD1c^+* dendritic cells (yellow dashed line), *COL1A1^+* fibrotic MΦ (purple dashed line), *MERTK^+LYVE1^+* fibrotic chondrocytes (green dashed line), *LTB^+CD69^+* myeloid cells (blue dashed line), and *S100A8^+S100A9^+* monocytes (pink dashed line). **(B)** Heatmap of normalized expression of differentially expressed genes among myeloid cell subsets.

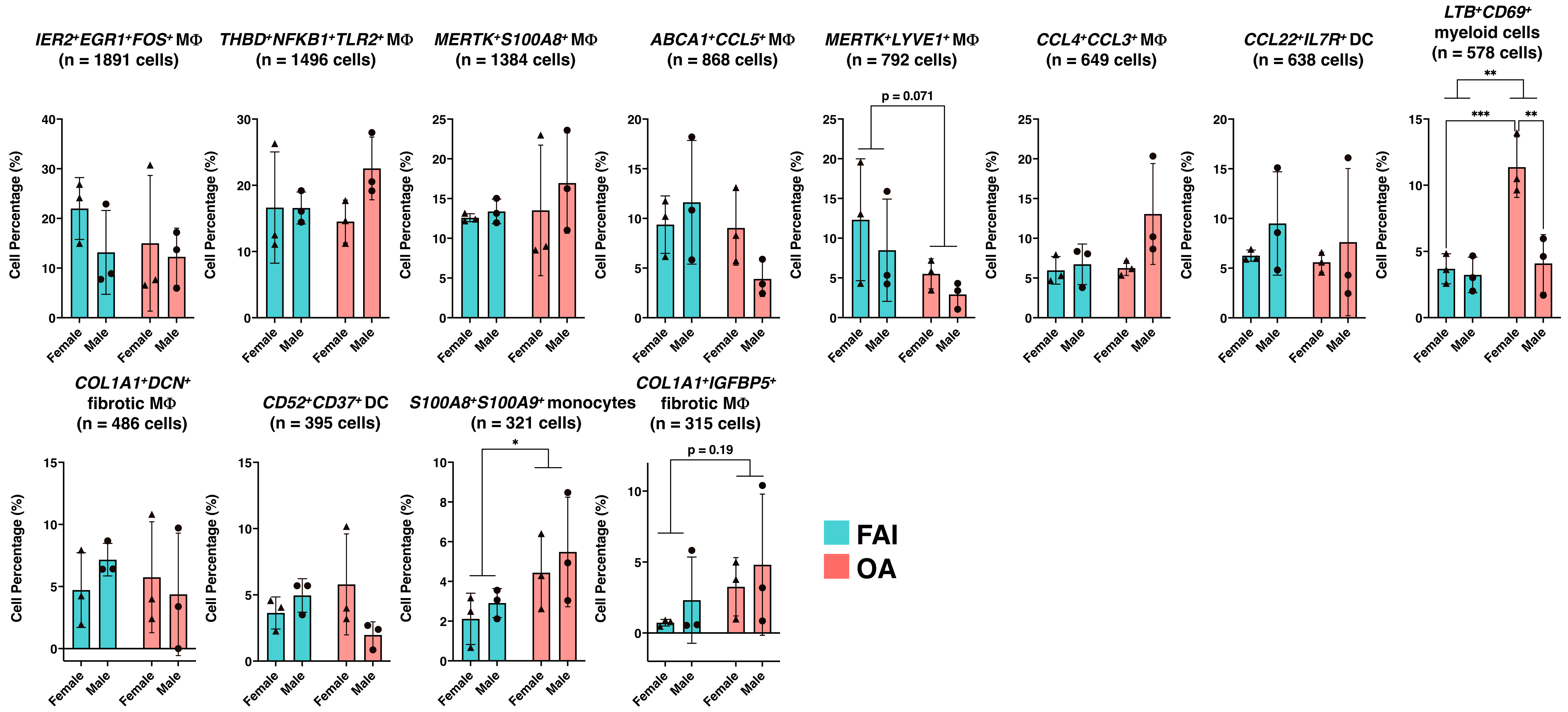


**Supplemental Fig. 6 | Altered percentages of *CD45+CD14+* synovial myeloid cells populations between FAI and OA patients.** Mean±SD. Two-way ANOVA with Fischer’s LSD test, * $p < 0.05$, ** $p < 0.01$.

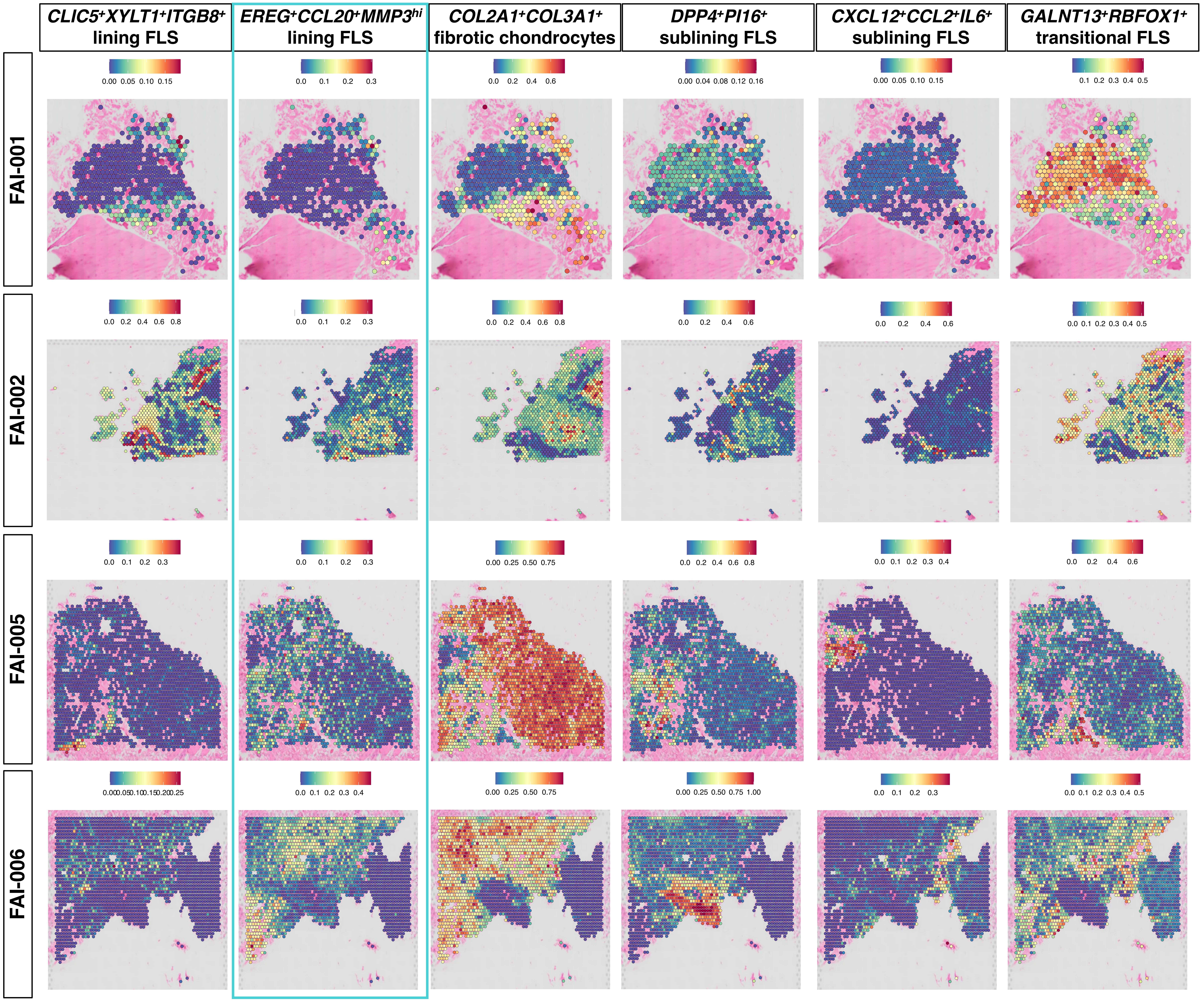

CLIC5+XYLT1+ITGB8+ lining FLS
EREG+CCL20+MMP3hi lining FLS
COL2A1+COL3A1+ fibrotic chondrocytes
DPP4+PI16+ sublining FLS
CXCL12+CCL2+IL6+ sublining FLS
GALNT13+RBFOX1+ transitional FLS
FAI-001
FAI-002
FAI-005
FAI-006

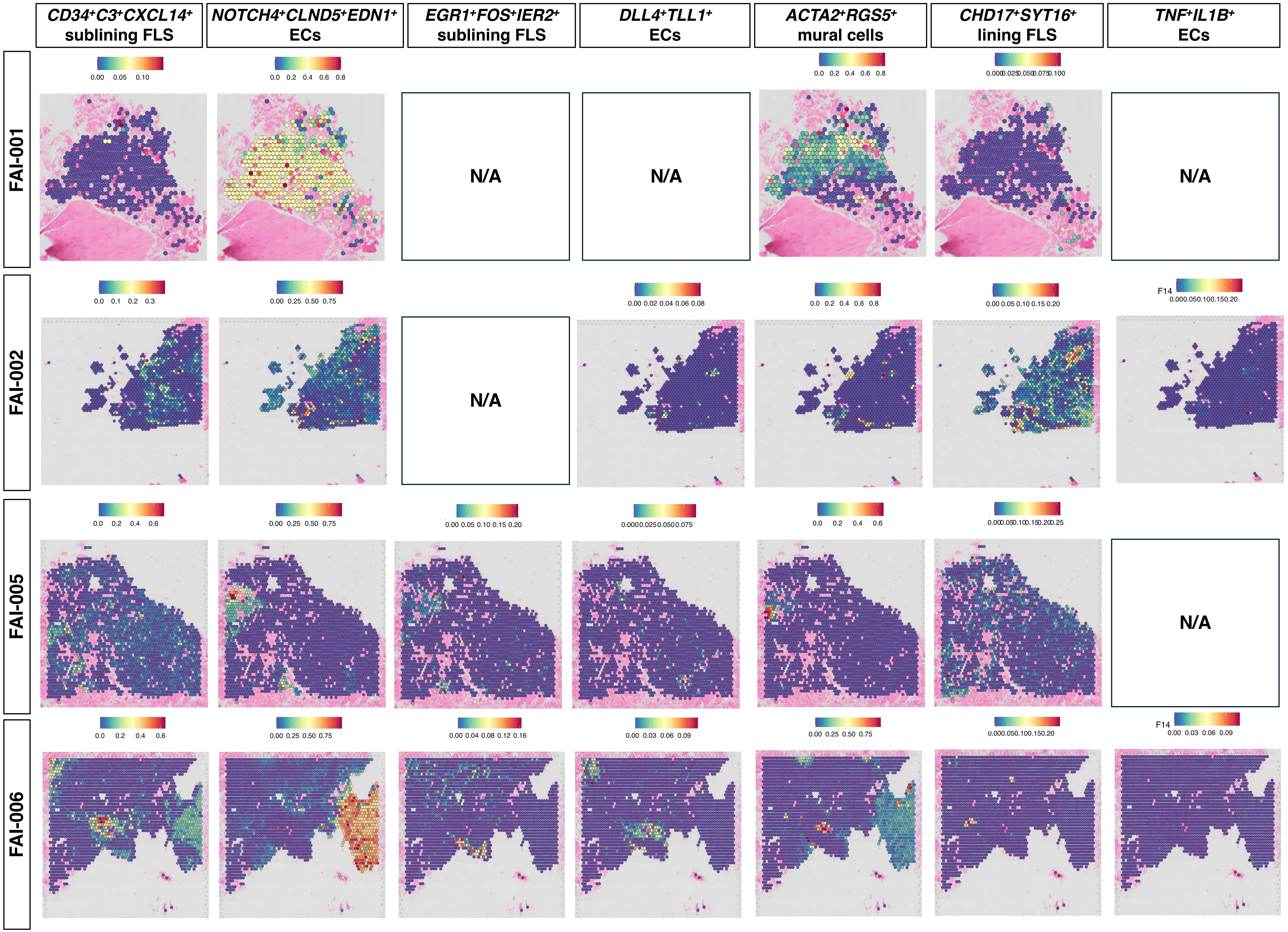


**Supplemental Fig. 7 | Spatial Feature Plots** of finer-grained clusters of *CD45*- non-hematopoietic cells in FAI synovium mapped to Spatial-seq spots. Note warmer color represents higher probability of that populations mapping to spatial-seq datasets and N/A means cluster was not mapped. This is likely due to the gene expression of DEGs for that cluster was masked by the gene expression of other clusters. Note population of *EREG+CCL20+MMP3hi* lining FLS in FAI synovium is highlighted in turquoise.

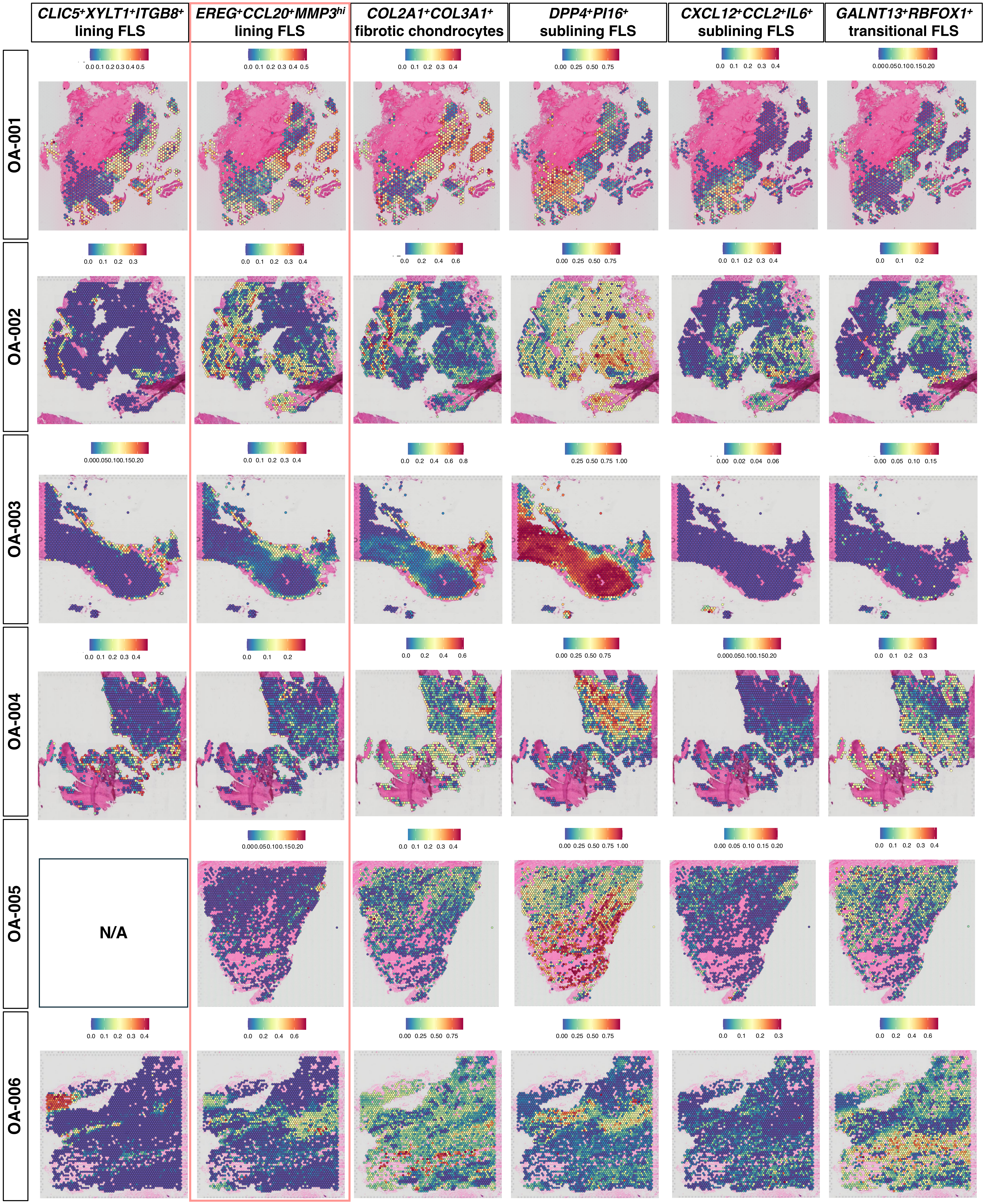

CLIC5+XYLT1+ITGB8+ lining FLS
EREG+CCL20+MMP3hi lining FLS
COL2A1+COL3A1+ fibrotic chondrocytes
DPP4+PI16+ sublining FLS
CXCL12+CCL2+IL6+ sublining FLS
GALNT13+RBFOX1+ transitional FLS
OA-001
OA-002
OA-003
OA-004
OA-005
OA-006
N/A

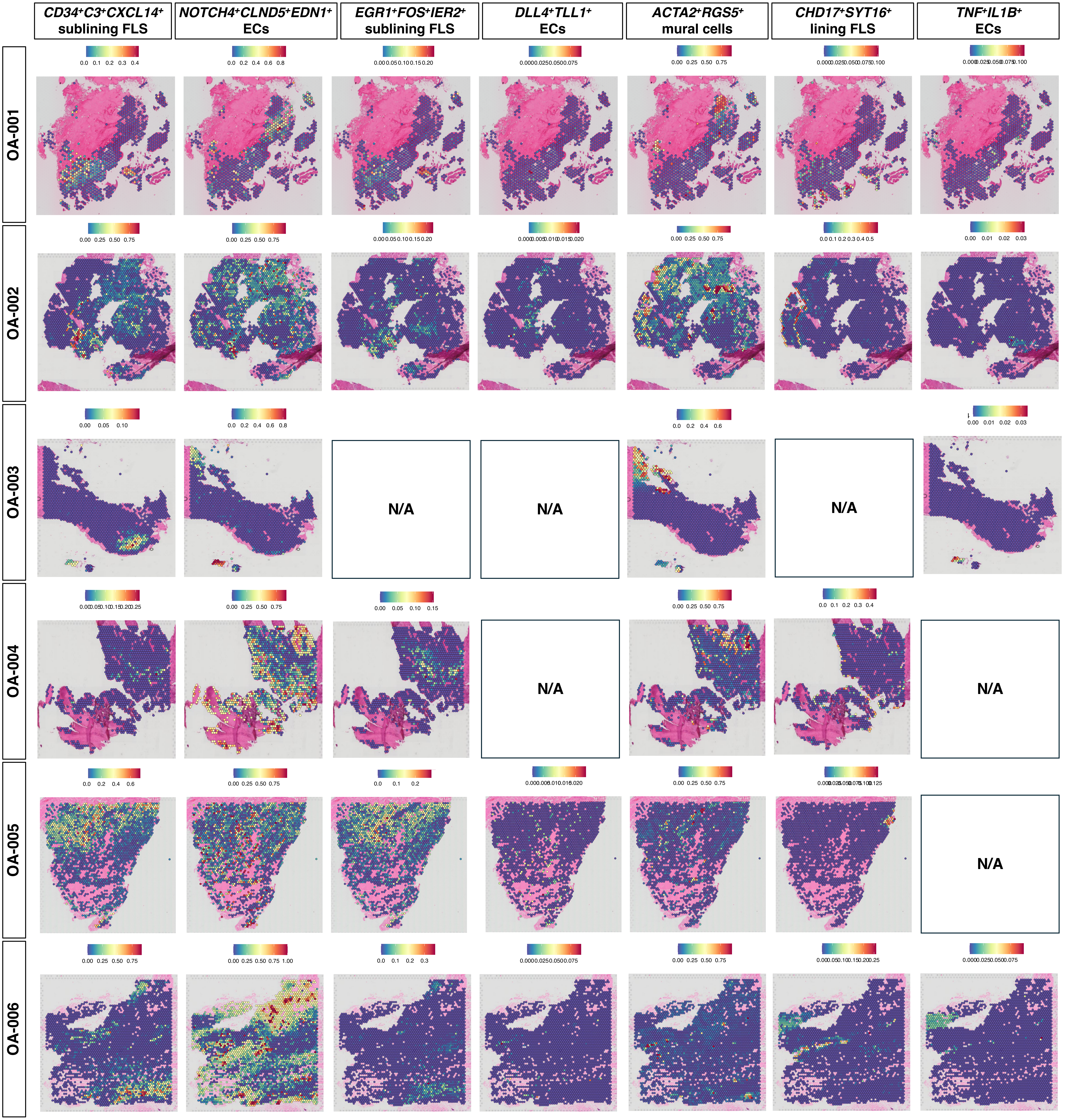


**Supplemental Fig. 8 | Spatial Feature Plots** of finer-grained cluster of *CD45*$^-$ non-hematopoietic cells in hip OA synovium mapped to Spatial-seq spots. Note warmer color represents higher probability of that populations mapping to spatial-seq datasets and N/A means cluster was not mapped. This is likely due to the gene expression of DEGs for that cluster was masked by the gene expression of other clusters. Note population of *EREG*$^+$*CCL20*$^+$*MMP3*$^{hi}$ lining FLS in FAI synovium is highlighted in salmon.

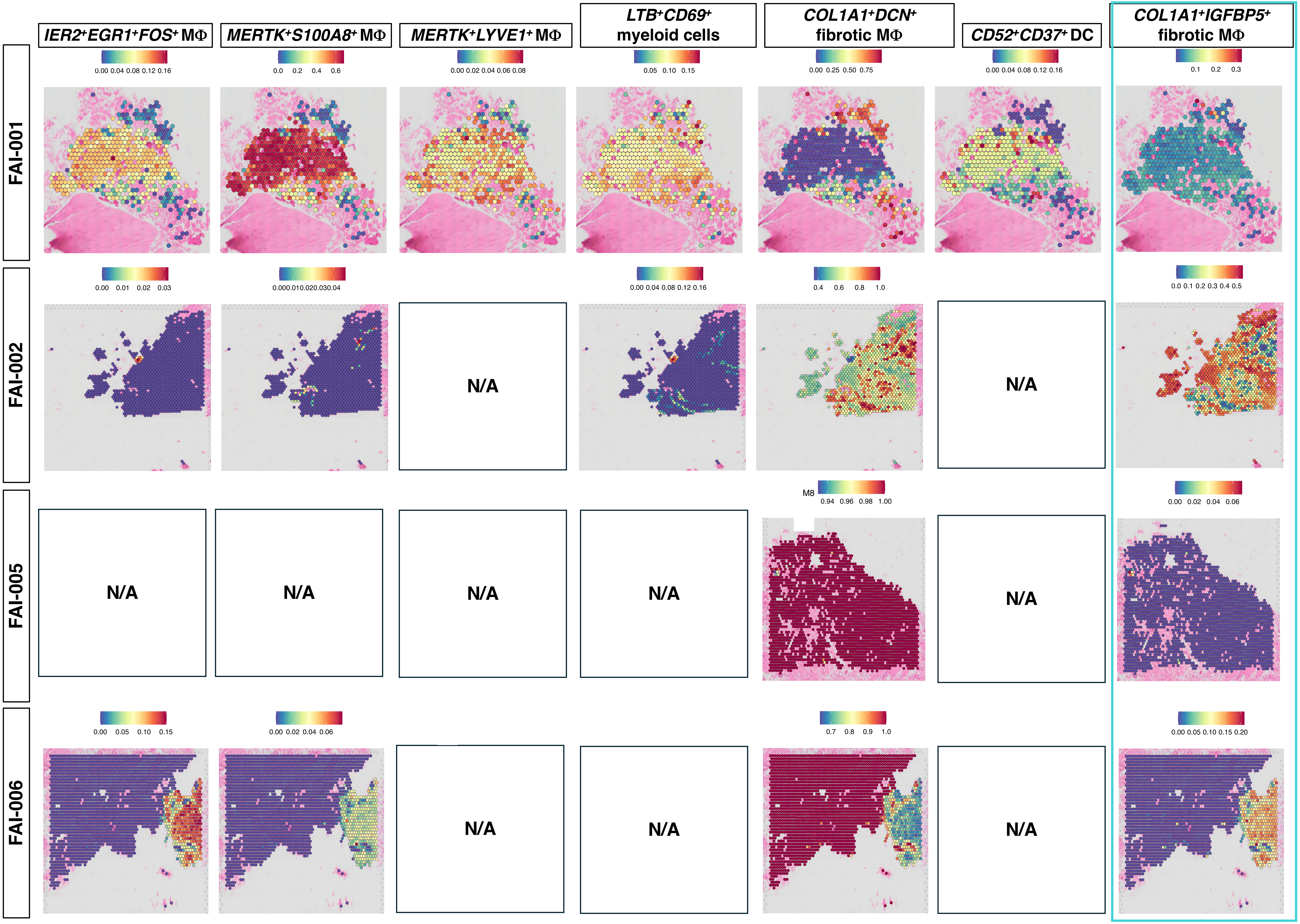


**Supplemental Fig. 9 | Spatial Feature Plots** of finer-grained cluster of *CD45⁺CD14⁺* myeloid cells in FAI synovium mapped to Spatial-seq spots. Note warmer color represents higher probability of that populations mapping to spatial-seq datasets and N/A means cluster was not mapped. This is likely due to the gene expression of DEGs for that cluster was masked by the gene expression of other clusters. Note population of *COL1A1⁺IGFBP5⁺* fibrotic MΦ in FAI synovium is highlighted in turquoise.

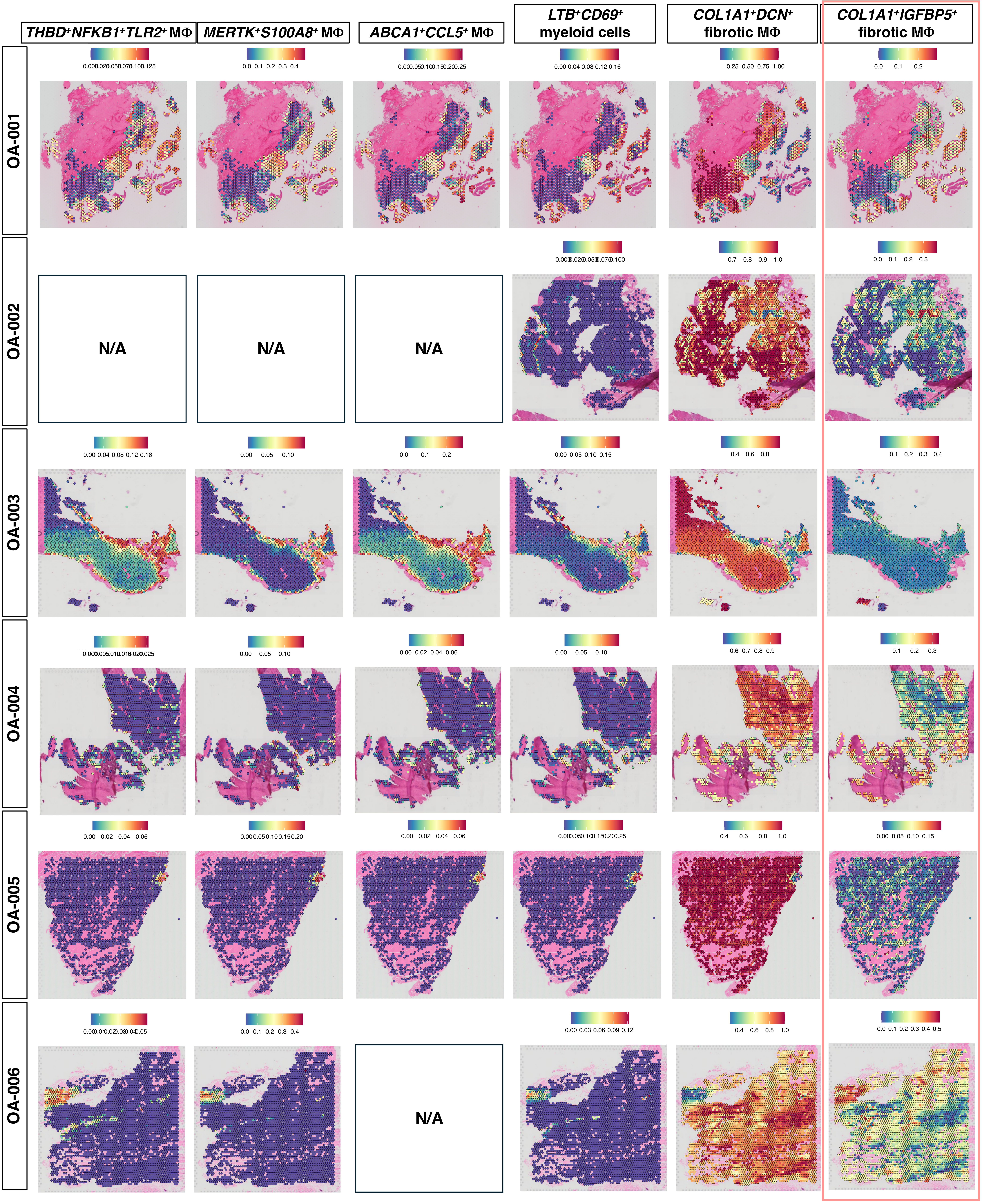


**Supplemental Fig. 10 | Spatial Feature Plots** of finer-grained cluster of *CD45+CD14+* myeloid cells mapped in hip OA synovium to Spatial-seq spots. Note warmer color represents higher probability of that populations mapping to spatial-seq datasets and N/A means cluster was not mapped. This is likely due to the gene expression of DEGs for that cluster was masked by the gene expression of other clusters. Note population of *COL1A1+IGFBP5+* fibrotic MΦ in OA synovium is highlighted in salmon.

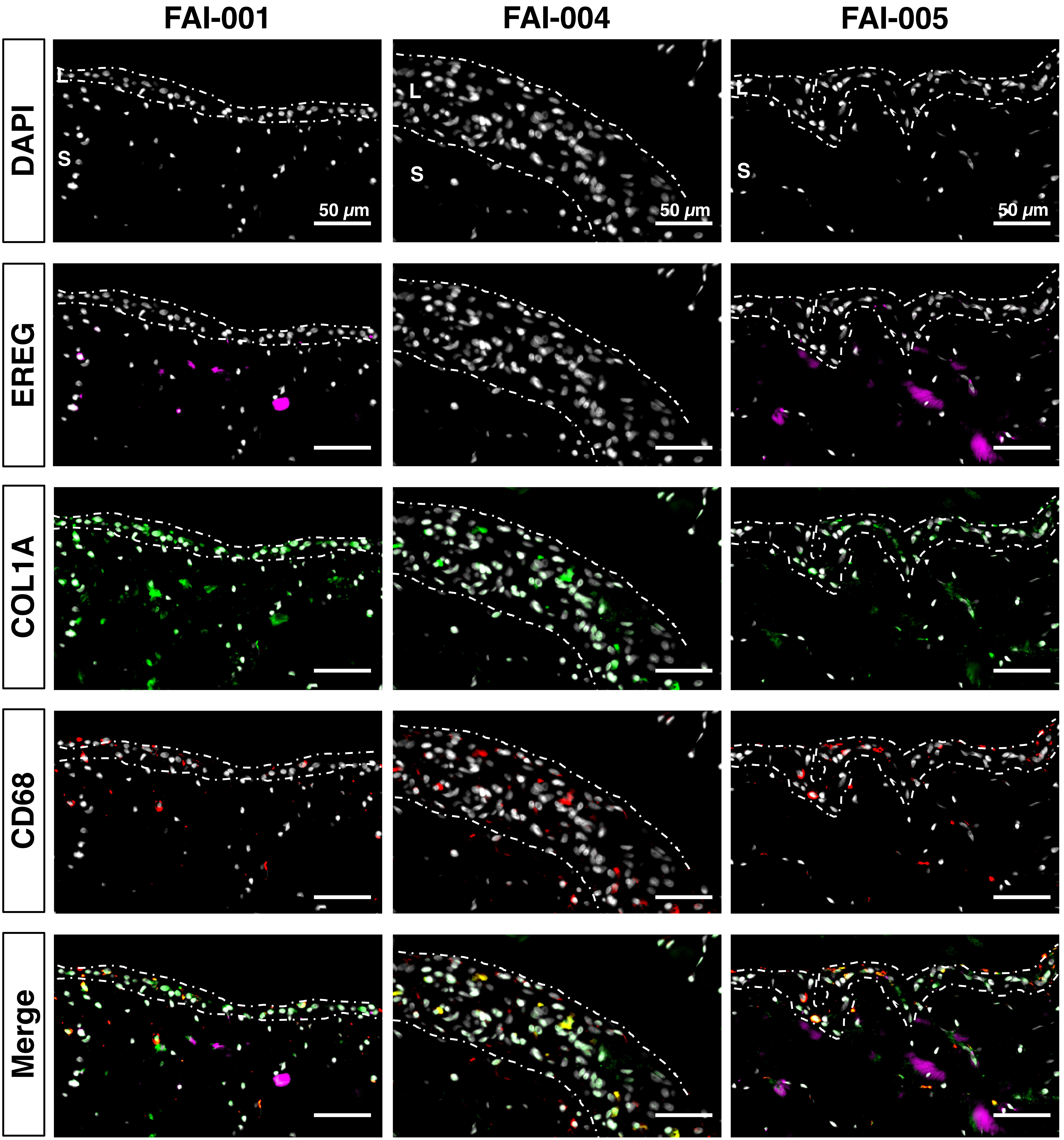


**Supplemental Fig. 11 | Immunofluorescent staining for epiregulin (EREG), collagen I (COL1) and CD68** to label *EREG*$^{+}$ and *CD68*$^{+}$ *COL1A1*$^{+}$ macrophages in FAI synovium (n=3).

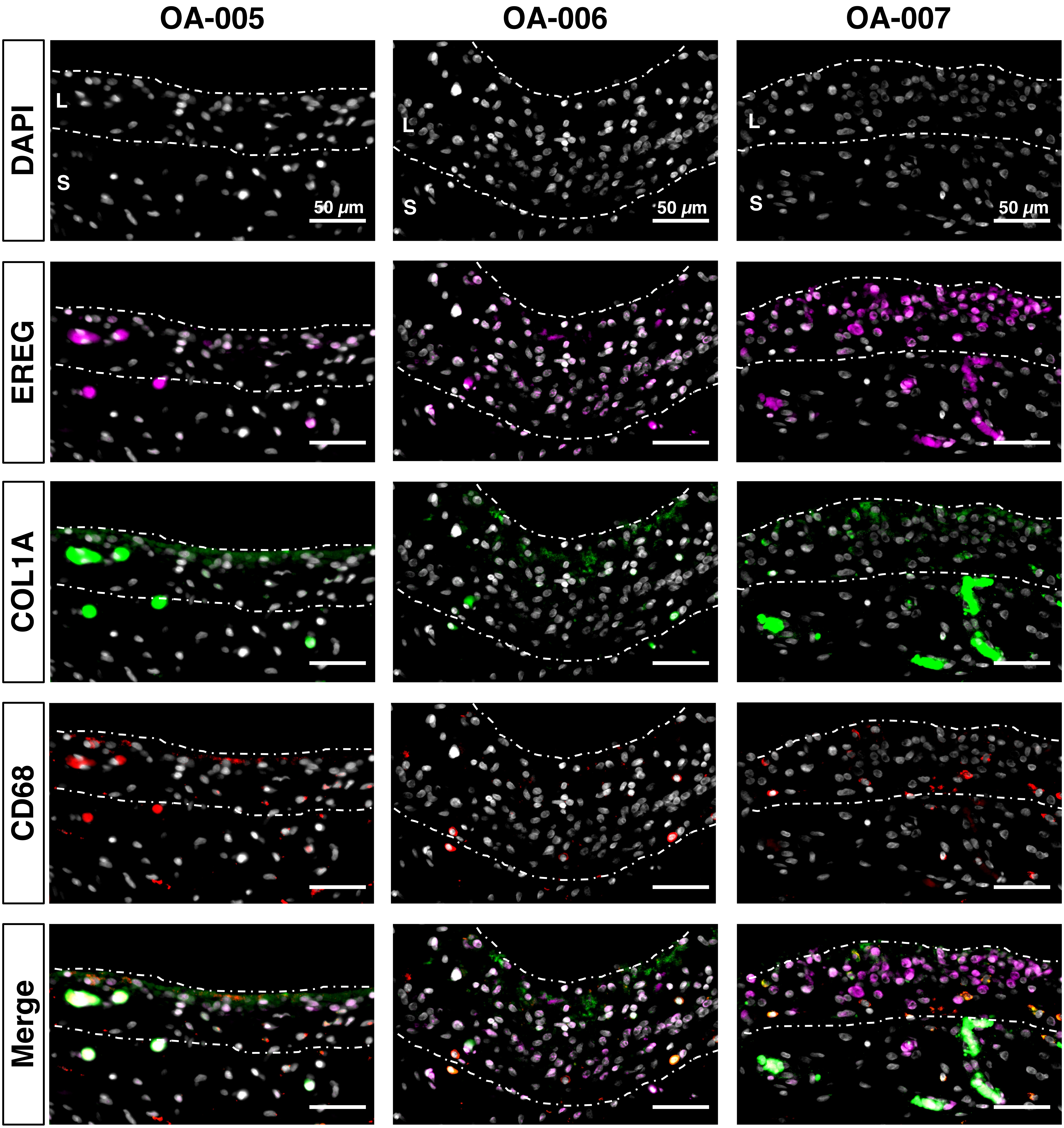


**Supplemental Fig. 12 | Immunofluorescent staining for epiregulin (EREG), collagen I (COL1) and CD68** to label *EREG*+ and *CD68*+ *COL1A1*+ macrophages in hip OA synovium (n=3).

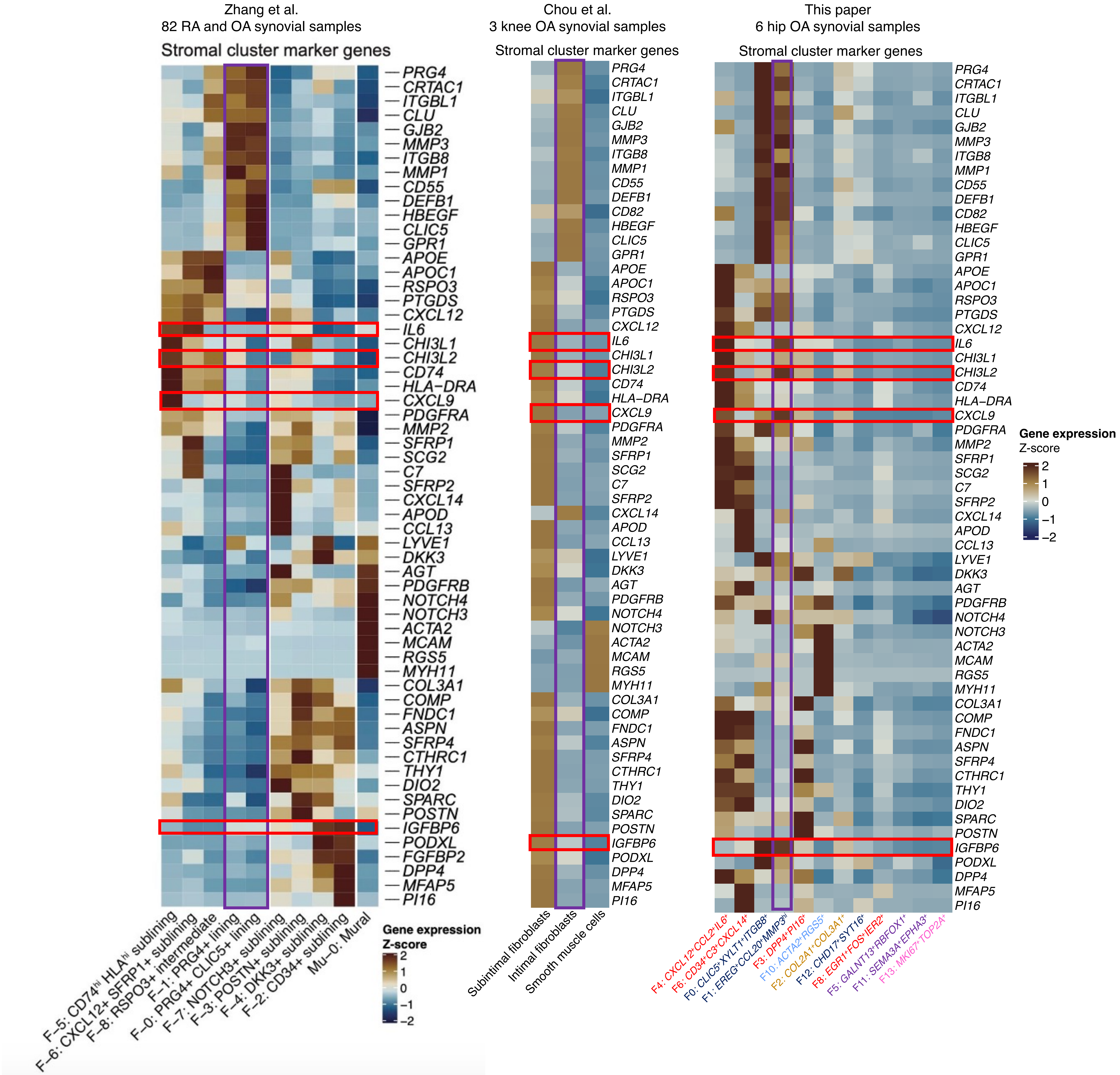


**Supplemental Fig. 13 | Heatmap comparison** of our datasets with publicly available datasets of stromal cells (fibroblasts and mural cells) in human RA/OA and knee OA synovia. Heatmap of human knee RA/OA synovium was adapted from Extended figure 7 in Zhang et al.. Note clusters of stromal cells in Chou et al. underwent supervised clustering to keep the cluster annotation and grouping identical to the original paper. Furthermore, heatmaps from our (**right**) and knee OA (**middle**) datasets were customized to fit stromal cluster marker gene heatmap from Zhang et al. Note lining fibroblasts from Zhang et al. and Chou et al. (highlighted in purple boxes) express similar gene sets as lining FLS in our datasets.

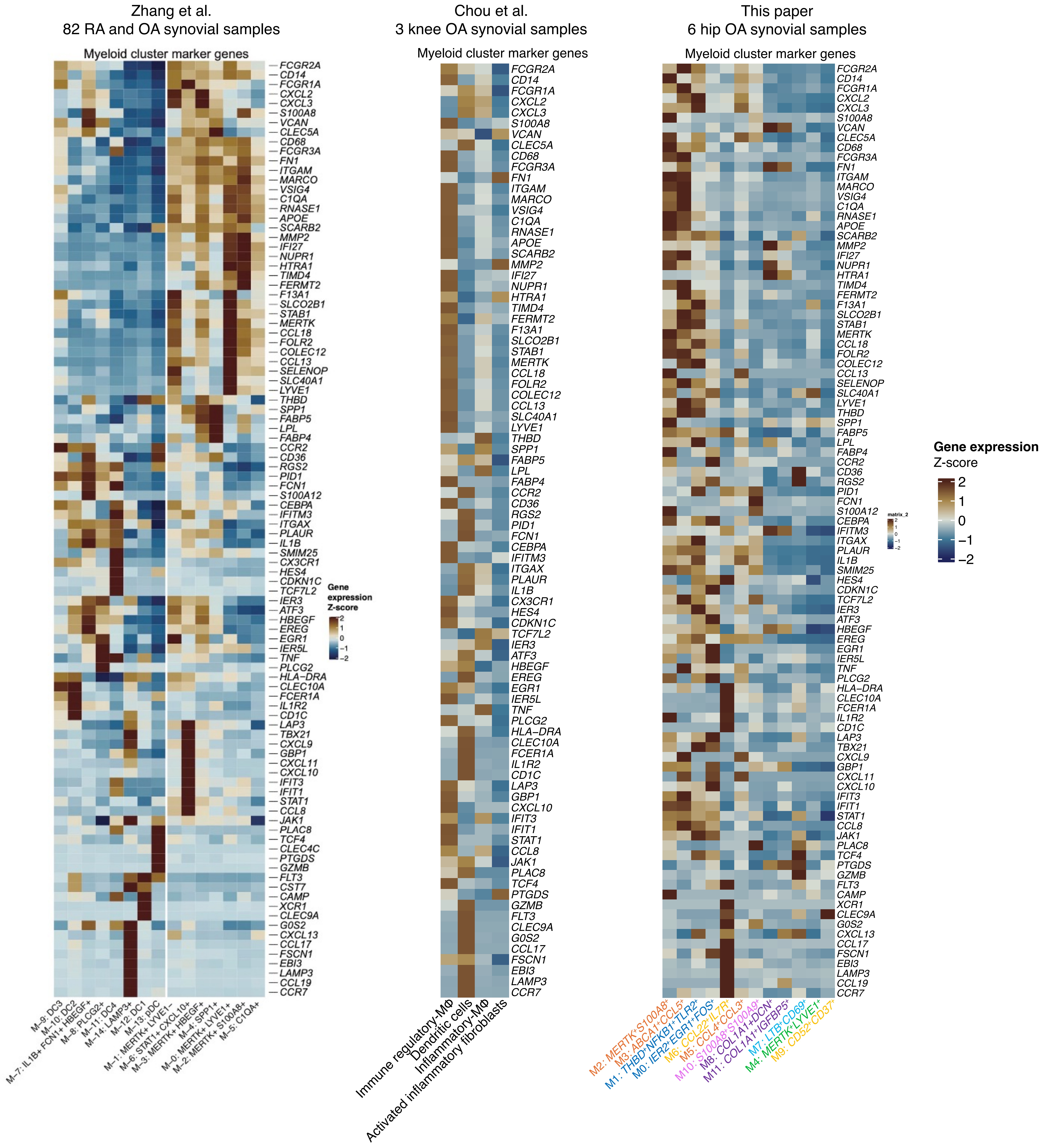


**Supplemental Fig. 14 | Heatmap comparison** of our datasets with publicly available datasets of myeloid cells in human RA/OA and knee OA synovia. Heatmap of human knee RA/OA synovium was adapted from Extended figure 6 in Zhang et al. Note clusters of myeloid cells in Chou et al. underwent supervised clustering to keep the cluster annotation and grouping identical to the original paper. Furthermore, heatmaps from our (**right**) and knee OA (**middle**) datasets were customized to fit myeloid cluster marker gene heatmap from Zhang et al.
